# Rapid spread of a symbiotic virus in a major crop pest following wide-scale adoption of Bt-cotton in China

**DOI:** 10.1101/2021.02.08.430243

**Authors:** Yutao Xiao, Wenjing Li, Xianming Yang, Pengjun Xu, Minghui Jin, He Yuan, Weigang Zheng, Mario Soberón, Alejandra Bravo, Kenneth Wilson, Kongming Wu

## Abstract

*Bacillus thuringiensis* (Bt) crops have been widely planted and the effects of Bt-crops on populations of the target and non-target insect pests were well studied. However, the effects of Bt-crops exposure on microorganisms that interact with crop pests haven’t previously been quantified. Here, we use laboratory and field data to show that infection of *Helicoverpa armigera* with a symbiotic densovirus (HaDV2) is associated with its enhanced growth and resistance to Bt-cotton. Moreover, field monitoring showed a much higher incidence of cotton bollworm infection with HaDV2 in regions cultivated with Bt-cotton than in regions without it, with the rate of densovirus infection increasing with increasing use of Bt-cotton. RNA-seq suggested resistance to both baculovirus and Cry1Ac were enhanced via the immune-related pathways. These suggest that the exposure to Bt-crops has selected for beneficial interactions between the target pest and a symbiotic microorganism that enhances its performance on Bt-crops under field conditions.

## Introduction

Transgenic crops expressing insecticidal Cry proteins from *Bacillus thuringiensis* bacteria, known as Bt-crops, have become important tools for the management of insect crop pests (*Carriere et al., 2003; Cattaneo et al., 2006; Hutchison et al., 2010; Shelton et al., 2002; Tabashnik et al., 2010*). Planting of Bt-crops effectively suppresses the targeted insects, decreasing insecticide use and promoting biocontrol services (*Bravo et al., 2011; Carriere et al., 2003; Cattaneo et al., 2006; Hutchison et al., 2010; Lu et al., 2010; Lu et al., 2012; Shelton et al., 2002; Tabashnik et al., 2002; Wu, 2010; Wu et al., 2008*). We have previously shown that commercialization of transgenic Bt-cotton in China brought significant changes in the ecology of insects utilizing the crop (*Lu et al., 2010 & 2012; Swiatkiewicz et al., 2014; Wu et al., 2008*). However, the effect of Bt-crops on other organisms such as symbiotic microbes, which could be playing important roles in the life cycle of insect populations, remains largely unknown.

Recently, we showed in laboratory trials that infection with a densovirus, *Helicoverpa armigera* densovirus-1 (HaDNV-1 / HaDV2) was associated with significantly enhanced resistance of cotton bollworm, *H. armigera*, to a baculovirus (*H. armigera* nucleopolyhedrovirus, HaNPV) (*Xu et al., 2014*), and there was some suggestion that the densovirus also increased resistance to Cry1Ac toxin in a Bt-susceptible strain of *H. armigera* (*Xu et al., 2014, 2017a and 2017b*). HaDV2 was found to be widespread in wild populations of *H. armigera* adults (>67% prevalence between 2008 and 2012) (*Xu et al., 2014*). The densovirus was mainly distributed in the fat body of the insect and could be both horizontally- and vertically-transmitted. Moreover, HaDV2-positive individuals showed faster development and higher fecundity than non-infected individuals. There was no evidence for a negative effect of HaDV2 infection on *H. armigera* in relation to other fitness-related traits, suggesting a possible mutualistic interaction between the cotton bollworm and HaDV2 (*Xu et al., 2014*).

Here, we further explore the interaction between *H. armigera*, HaDV2 and Bt-cotton, to establish its relevance to field populations and to test the hypothesis that the widespread adoption of Bt-cotton in China has selected for cotton bollworm carrying the symbiotic densovirus. Laboratory experiments show that HaDV2 infection in both Cry1Ac-resistant and Cry1Ac-susceptible strains of cotton bollworm enhances larval resistance to Bt and overall fitness. Field experiments indicate that *H. armigera* populations infected with HaDV2 also have higher resistance to Bt-cotton relative to those not infected with HaDV2. Moreover, field monitoring over a ten-year period indicates that the frequency of HaDV2-infected cotton bollworm is significantly higher in regions planted with Bt-cotton than in those areas where Bt-cotton is not grown, and that in Bt-cotton-growing areas the prevalence of HaDV2 infection increases with time since Bt-cotton adoption and with the proportion of cotton that is grown which is transgenic. Further, we found that increased Bt resistance is associated with activated immune pathways to HaDV2-infection. These results indicate that increased HaDV2 infection in cotton bollworm is correlated with the wide-scale adoption of Bt-cotton in China. Our data are consistent with the notion that exposure to Bt-crops selects for beneficial interactions between the target pest and a microorganism that enhances their fitness in response to Bt exposure under field conditions.

## Results

### HaDV2 infection increases Cry1Ac resistance level in *H. armigera*

Previous laboratory bioassays suggested that when a Cry1Ac-susceptible strain of *H. armigera* was infected with HaDV2 it increased its resistance to the Cry1Ac toxin (*Xu et al., 2014*). To explore the generality of this finding, we first analyzed the effect of HaDV2 infection in different *H. armigera* populations that differ in their susceptibility to Cry1Ac due to different mechanisms of resistance. Two susceptible *H. armigera* strains infected with HaDV2 (96S and LF) both showed 1.5 times greater resistance levels to Cry1Ac toxin, relative to their corresponding non-infected controls (*Supplementary table 1 and 2*). In the case of the Cry1Ac-resistant strains (BtR, 96CAD, LFC2, LF5, LF60, LF120 and LF240), infection with HaDV2 again showed a significant increase in their resistance levels relative to the corresponding strains without HaDV2 infection, ranging between 30% and 130% enhanced resistance (*Figure 1*). The slope of the regression line is significantly greater than one (t-test: t = 2.853, df = 6, P = 0.029), suggesting that the benefits of carrying HaDV2 may increase with increasing levels of Bt resistance. Logistic regression confirmed that for a given strain of *H. armigera*, larvae harboring HaDV2 were significantly more resistant to Cry1Ac (GLM: Strain: χ^2^_8_ = 36.57, P < 0.0001, log10 (Cry1Ac toxin concentration): χ^2^_1_ = 848.16, P < 0.0001; Strain* log10(Cry1Ac toxin concentration): χ^2^_8_ = 97.30, P < 0.0001; HaDV2-status: χ^2^_1_ = 9.53, P = 0.0020). However, there was no evidence that the benefits of hosting HaDV2 are affected by the resistance level of the *H. armigera* strain, as reflected in the LC50 of the non-infected insects (χ^2^_1_ = 0.06, P = 0.80).

**Figure 1.**
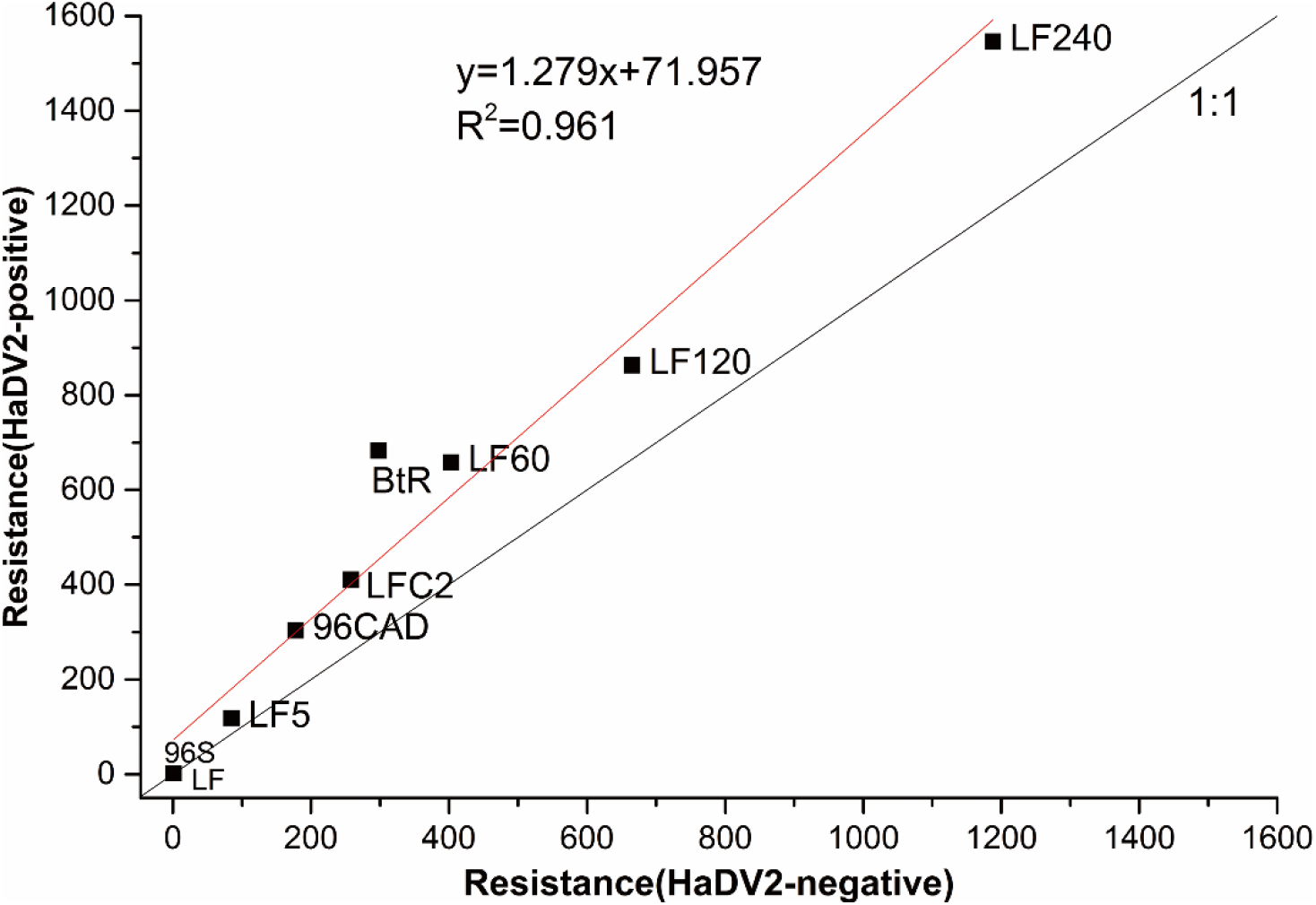
Relationship of different *H. armigera* strains’ resistance levels with or without HaDV2 infection. The x-axis is the resistance levels of different strains (LF, 96S, LF5, LF60, LF120, LF240, LFC2, 96CAD and BtR) without HaDV2 infection (HaDV2-negative); the y-axis is the resistance levels of different strains (LF, 96S, LF5, LF60, LF120, LF240, LFC2, 96CAD and BtR) with HaDV2 infection (HaDV2-positive). In both cases, the resistance level is the ratio of LC_50_ for the focal strain relative to the LC_50_ of the HaDV2-negative 96S strain larvae. The regression line is described by the following equation: y = 1.279x + 71.957, R²= 0.961, F =170.7, df = 1,7, P < 0.0001).

### HaDV2 infection reduces the fitness cost of H. armigera associated to Cry1Ac-resistance evolution

To determine whether infection with HaDV2 reduces the costs associated with evolving resistance to Bt, a range of fitness traits (*Supplementary table 3*) were measured in four strains of *H. armigera* that have different Bt-resistance levels (LF, LF5, LF60 and LF240) and were infected or not infected with HaDV2. Enhanced Bt-resistance in *H. armigera* LF, LF5, LF60 and LF240 strains was associated with lower larval survival rates, prolonged larval and pupal development (in both sexes), reduced pupal weight, lower adult emergence rate and reduced fecundity and egg hatch rate; in contrast, sex ratio at emergence and the longevity of adults of both sexes were not influenced by the capacity to resist Cry1Ac toxin (*Supplementary table 3 and 4; Supplementary figure 1*). When traits were combined to estimate the fundamental net reproductive rates of the four *H. armigera* strains, *R*_0_ (*Wang et al., 2016*), this revealed that in the absence of HaDV2 infection, the fitness of the most Bt-resistant strain (LF240) was around 40% of that of the most Bt-susceptible strain (LF), consistent with a large fitness cost of resistance (*Figure 2*); strains with intermediate levels of resistance (LF5, LF60) suffered a lower cost of resistance (*c*. 30% reduction in fitness).

**Figure 2.**
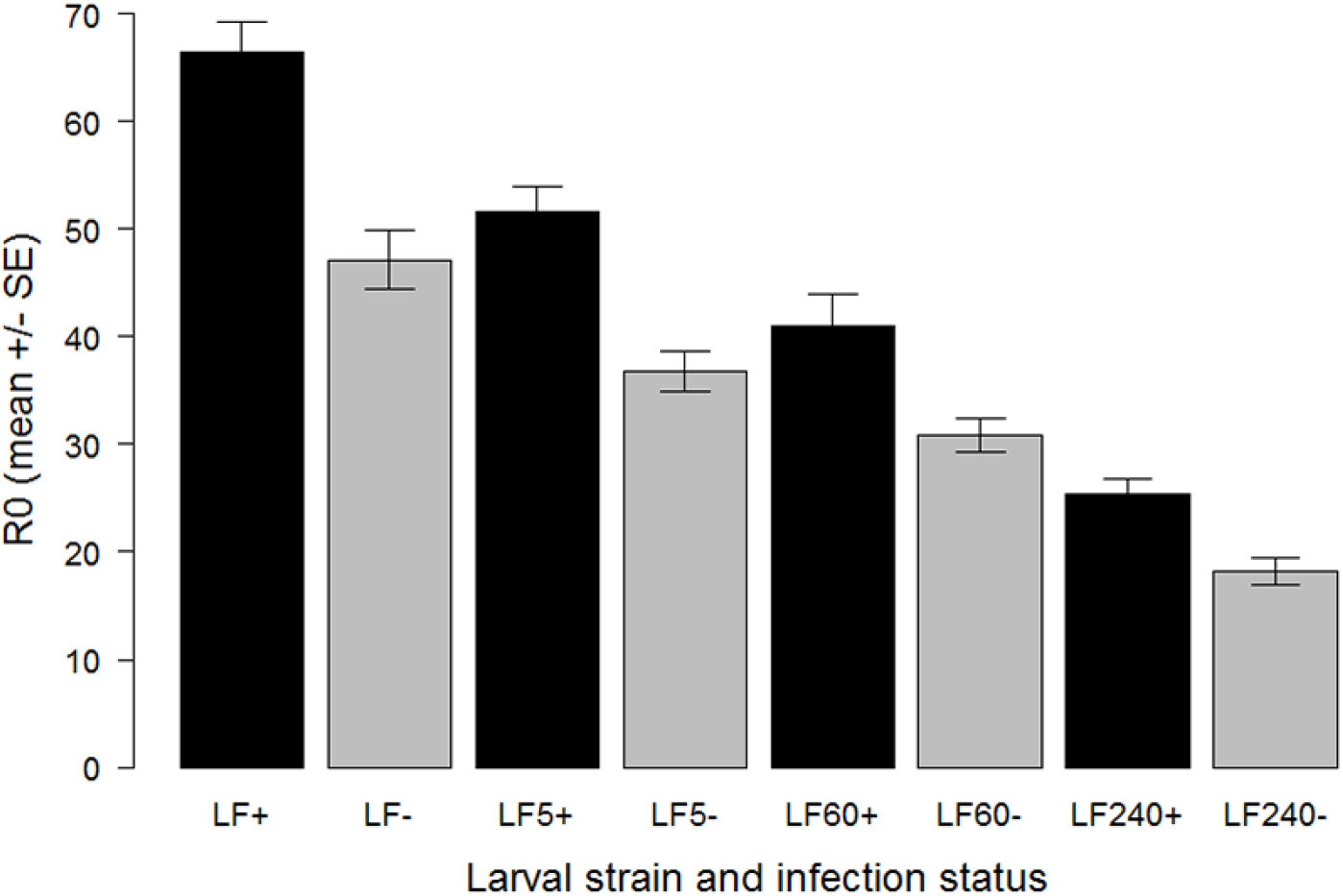
Effects of HaDV2 infection on the net reproductive rate (R_0_) in four *H. armigera* strains differing in their resistance to Bt and not exposed to Cry1Ac toxin. Mean R_0_ is calculated as the number of female offspring per female that reaches adulthood. The error bars are bootstrapped standard errors.

Infection with HaDV2 rescued, or partially rescued, this fitness loss, in all four *H. armigera* strains showing a significant increase in *R*_0_ relative to their non-infected counterparts, averaging around 38% higher (paired t-test: t = 4.831, df = 3, P = 0.017; *Figure 2*). Indeed, when infected with HaDV2, the *R*_0_ values of two of the three resistant strains (LF5 and LF60) were comparable to that of the non-infected susceptible (LF) strain (*Figure 2*).

To quantify the fitness cost associated with Cry1Ac-resistance in the more realistic context of multiple plant defenses, the growth rate of the different strains of *H. armigera* larvae was also analyzed on Bt-cotton plant leaves. Larval weight after nine days growth was significantly affected by the cotton variety (Bt or non-Bt) and by the HaDV2 infection-status (*Supplementary table 5 and 6*), with larvae generally being heavier when fed with non-Bt-cotton than with Bt-cotton, and heavier for HaDV2-infected larvae than for larvae not infected with HaDV2; larvae were also heavier when they expressed lower levels of Bt-resistance, indicating that the cost of resistance is reflected in larval growth (*Supplementary table 5 and 6*). None of the interactions between these three main effects explained any additional variation (model comparison with and without interaction terms: F = 0.601, df = 14, P = 0.86), suggesting that the effects of host plant, *H. armigera* strain and infection-status on larval growth were additive.

### HaDV2 infection levels in field populations of H. armigera have increased over the adoption period of Bt-cotton

To determine if infection with HaDV2 could increase the performance of *H. armigera* when exposed to Bt-cotton in the field, we collected *H. armigera* moths from Xiajin (Shandong province) and Anci (Hebei province) in northern China, two locations where Bt-cotton has been widely planted over the last decade (*An et al., 2015*). Across two successive years, the prevalence of HaDV2 was extremely high (98% of 637 larvae in 2015; 97% of 180 larvae in 2016). Moreover, across both years, the relative average development rates (RADR) (*An et al., 2015*) of individuals infected with HaDV2 were significantly higher than that of larvae not infected with the virus (0.62 *vs* 0.52 in 2015; 0.61 *vs* 0.53 in 2016) (linear model: infection status: F = 28.80; df = 1,815, P < 0.0001; year: F = 2.10, df = 1,814, P = 0.15; infection*year: F = 0.57, df = 1,813, P = 0.57) (*Figure 3*).

**Figure 3.**
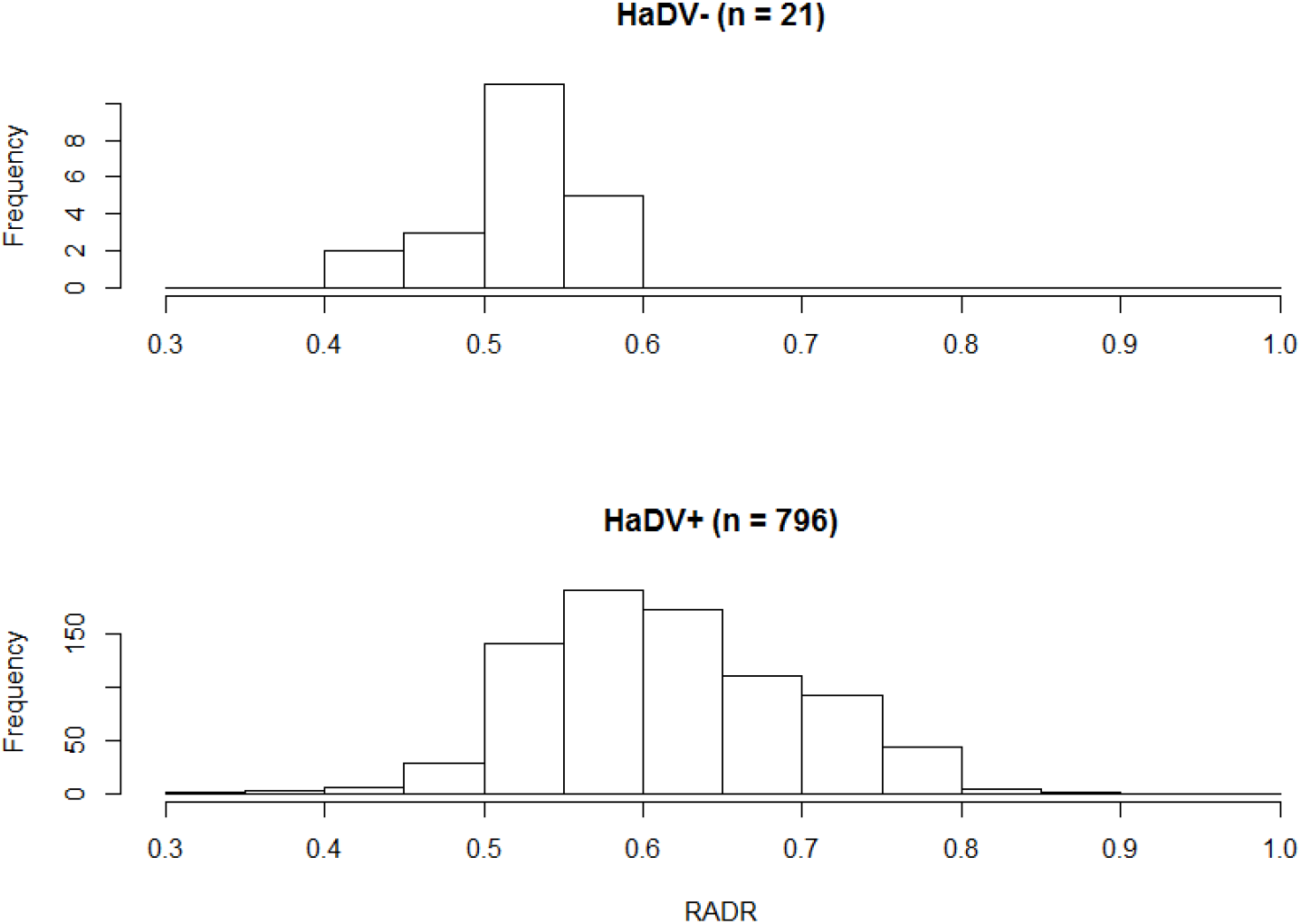
Frequency distributions for RADR scores for HaDV2 positive and negative insects. The data were collected from field-collected insects from Xiajin and Anci in 2015 and 2016.

Given the apparent selective advantage of HaDV2 infection for insects feeding on Bt-cotton, we predicted that over time we would observe an increase in HaDV2 infection rates in the field and that this would be associated with a temporal increase in average development rates for larvae feeding on Bt-cotton plants. As predicted, over the ten-year period between 2007 and 2016, at both Xiajin and Anci provinces, HaDV2 infection rates increased significantly over time (logistic regression: Xiajin: χ^2^_1_ = 405.79, P < 0.0001; Anci: χ^2^_1_ = 325.21, P < 0.0001) (*Figure 4A and 4B*). Associated with this, there was a significant temporal increase in larval development rates (RADR) at both locations (linear models: Xiajin: F = 5.474, df = 1,8, P = 0.047; Anci: F =23.256, df = 1,8, P = 0.0047) (*Figure 4C and 4D*). Moreover, across the ten years at both monitoring locations, there was a strong positive association between HaDV2 infection levels and RADRs, consistent with a possible causal relationship between these two temporal trends (linear models: Xiajin: F =23.826, df = 1,9, P = 0.001; Anci: F =13.676, df = 1,9, P = 0.006) (*Figure 4E and 4F*).

**Figure 4.**
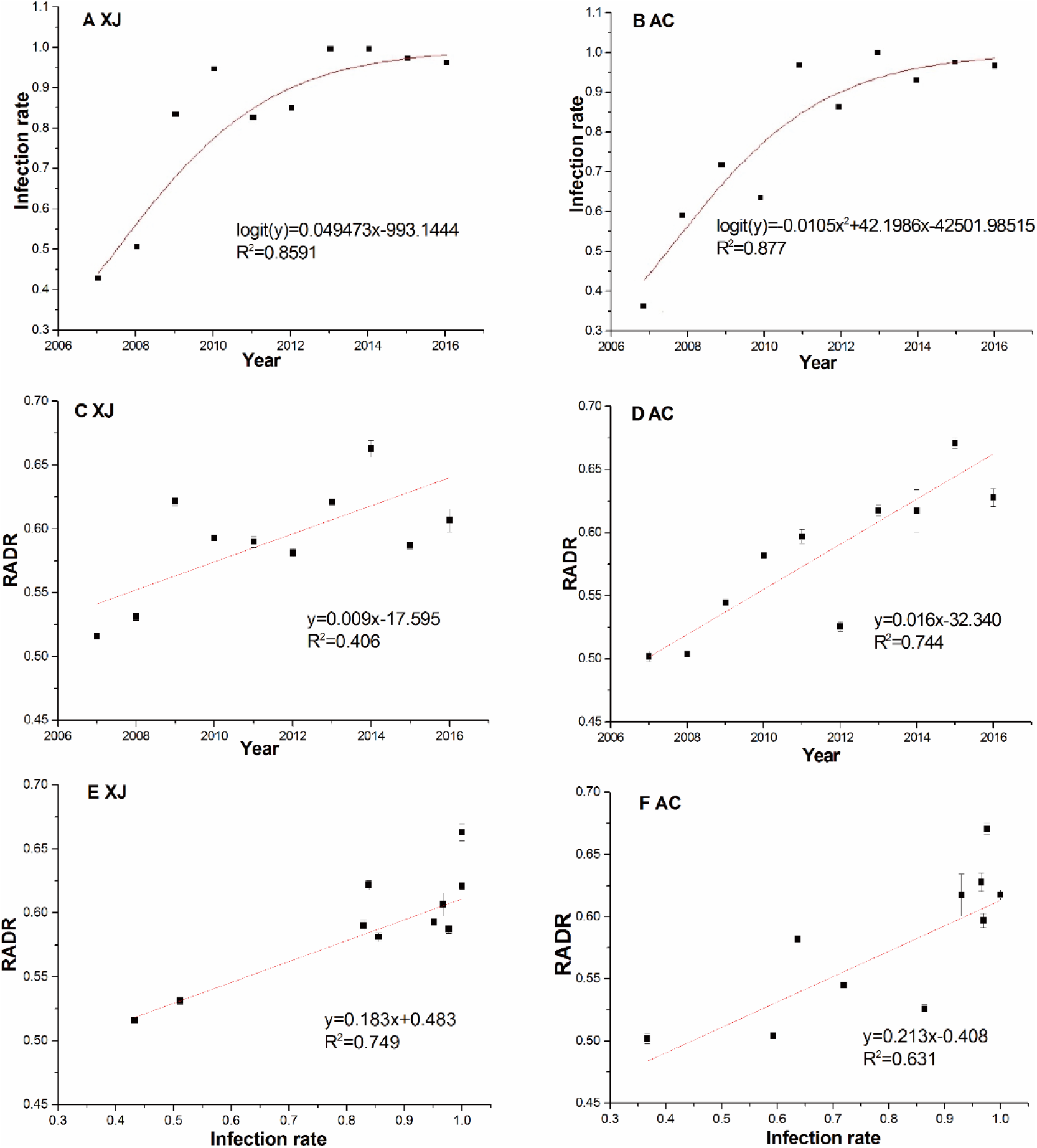
HaDV2 infection rate and RADR dynamics and their relationship for each year in the Xiajin and Anci populations during 2007–2016. (A) Relation between HaDV2 infection rate of larvae in Xiajin populations and planting year of Bt-cotton. Logistic regression model of HaDV2 infection rate, logit (y) = 0.49473x-993.1444, R² = 0.8591, χ^2^ = 405.79, df = 1, P < 0.0001. (B) Relation between HaDV2 infection rate of larvae in Anci populations and planting year of Bt-cotton. Logistic regression model of HaDV2 infection rate, logit (y) = −0.0105x^2^+42.1986x – 42501.98515, R² = 0.877, χ^2^ = 325.21, df = 1, P < 0.0001.(C) Relation between RADR of larvae in Xiajin populations and planting year of Bt-cotton. Linear model of RADR, y = 0.009x – 17.595, R²= 0.406, F =5.474, df = 1, 8, P = 0.047. (D) Relation between RADR of larvae in Anci populations and planting year of Bt-cotton. Linear model of RADR, y = 0.016x – 32.340, R^2^ = 0.744,F = 23.256, df = 1,8, P = 0.001. (E): Relationship of larvae RADR in Xiajin population and HaDV2 infection rate during the years 2007-2016, each data point is a different year, in the Linear model of RADR, y = 0.183x +0.438, R²= 0.749, F = 23.826, df = 1, 9, P = 0.001; (F): Relationship of larvae RADR in Anci populations and HaDV2 infection rate during the years 2007-2016, each data point is a different year, in the Linear model of RADR, y = 0.213+ 0.408R²= 0.625, F = 13.676, df = 1,9, P = 0.006. The bars are the standard error of the mean RADR for the field-derived strains tested in each year.

### Across regions the HaDV2 infection rates increases with increasing exposure to Bt-cotton

To further test the association between *H. armigera* densovirus infection levels and the adoption of Bt-cotton, we monitored HaDV2 infection rates at 36 locations across sixteen provinces during the period 2014 – 2016, including locations planted with transgenic Bt-cotton (29 monitoring points across 12 provinces) and locations where Bt-cotton has not been planted (9 monitoring points across 4 provinces) (*Supplementary figure 2; Supplementary table 7*). Across all three years, HaDV2 infection levels in *H. armigera* were significantly higher at locations where Bt-cotton was planted (mean = 82%) than in those where it was not (15%) (*Supplementary figure 3*) (logistic regression: crop (Bt vs non-Bt): χ^2^_1_ = 354.15, P < 0.0001). There was also a significant year-by-crop interaction (χ^2^_1_ = 24.13, P < 0.0001) due to HaDV2 infection levels being uniformly high across the three years at sites growing Bt-cotton (81-90%), whereas infection levels gradually increased from 2014 to 2016 at non-Bt sites (12%, 16% and 44%, respectively).

Moreover, across the provinces where Bt-cotton is grown, the mean prevalence of HaDV2 in *H. armigera* between 2014 and 2016 increased with the number of years since Bt-cotton was first introduced (χ^2^_1_ = 173.59, P < 0.0001) (*Figure 5a and Supplementary figure 3*) and also increased as the proportion of cotton that was transgenic increased (χ^2^_1_ = 5.34, P = 0.021) (*Figure 5b*). HaDV2 prevalence was not correlated with the proportional area of any of the other crops grown (χ^2^_1_ < 1.310, P > 0.25). Adding environmental variables to this minimal model, either singly or in combination, did not significantly improve the model fit (average rainfall: χ^2^_1_ = 0.110, P = 0.74; average temperature: χ^2^_1_ = 0.155, P = 0.69; average altitude: χ^2^_1_ = 0.001, P = 0.98; rainfall + temperature + altitude: χ^2^_3_ = 0.572, P = 0.90). These results are consistent with the notion that the benefits of HaDV2 infection are greatest for *H. armigera* exposed to Cry1Ac-producing cotton and that selection favoring HaDV2 infection increases the longer the insects are exposed to the Bt-cotton.

**Figure 5.**
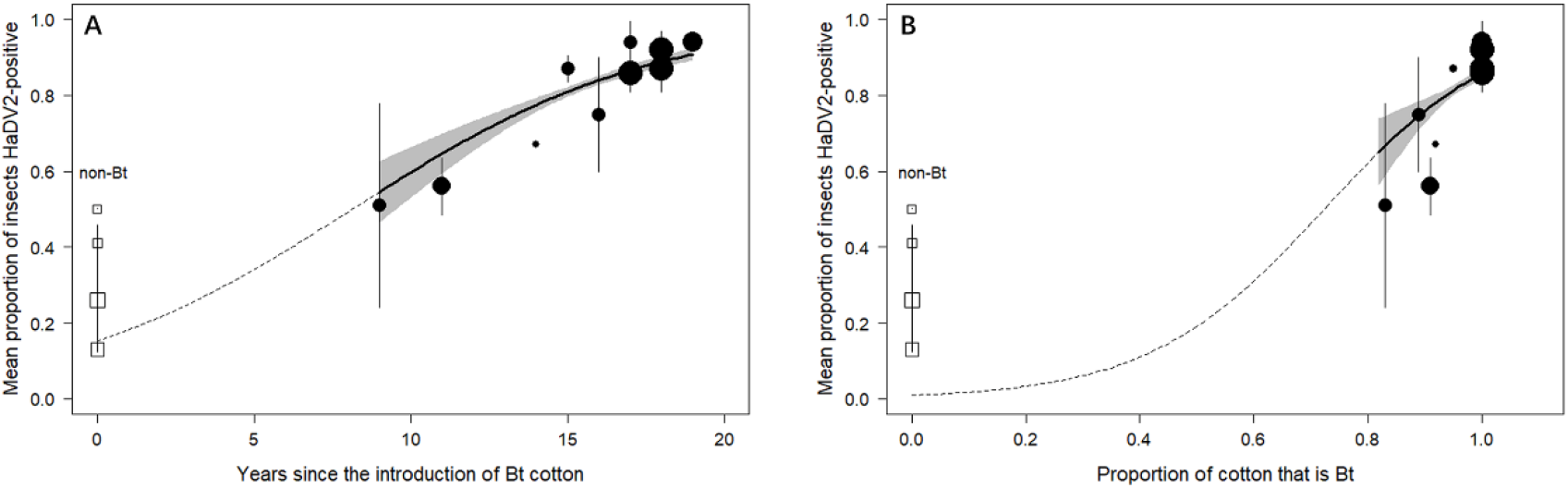
HaDV2 infection rate has increased significantly since Bt-cotton was first introduced and was positively related to the Bt adoption in all cotton areas. (A) Temporal changes in the infection rate of HaDV2 since the introduction of Bt-cotton. (B) Changes in the infection rate of HaDV2 according to the proportion of Bt cotton in all cotton. Each symbol represents an individual province sampled for densovirus over three years (2014-2016); the mean virus prevalence (+ standard error) over those three years is shown. Symbol size reflects sampling effort and represents data from >1500 insects. Circles represent the twelve provinces where Bt-cotton is grown; squares are the four provinces where Bt-cotton is not grown. The solid line represents the logistic regression (+ standard error, shaded zone) describing the relationship between virus prevalence and years since the introduction of Bt-cotton to a province for the twelve Bt-cotton-growing provinces only. The dashed line extrapolates this regression line to Year 0. The detailed information is summarized in supplementary table 7 and supplementary figure 2.

### HaDV2 infection activated the immune pathways in the cotton bollworm

To try to understand better the mechanisms increasing the Cry1Ac resistance levels and enhanced fitness of HaDV2-infected insects, we conducted an RNA sequencing experiment (*Supplementary table 8*).

The principal component analysis (PCA) of the transcriptome with DEG data clearly distinguished HaDV2-positive from -negative individuals at three different time points: 24 h, 48 h and less so at 72 h after HaDV2 inoculation (*Supplementary figure 4a, b, c*). Taken together with the hierarchical clustering of these DEGs, these results suggest that the HaDV2 has a major effect on the gene expression profiles of their hosts.

We performed pathway enrichment analysis on the DEGs, focusing particular attention on pathways related to the development and immune systems (*Supplementary figure 4d, e, f*). Genes in Jak-STAT immune signaling pathway, which are related to some antiviral and antibacterial mechanisms, were significantly enriched and up-regulated in the HaDV2-infected larvae at 24 h and 48 h (*Supplementary figure 4d, e, and 5a, b*), but not at 72 h (*Supplementary figure 4f*). Interestingly, genes in ABC transporters pathway at 48 h (*Supplementary figure 5c*), the mitogen-activated protein kinase (MAPK) signaling and lysosome pathways at 72 h (*Supplementary figure 5d, e*), which are related to antimicrobial immune response, are significantly enriched and up-regulated (*Supplementary figure 4e, f*). Genes in pathways related to development were also significantly enriched, e.g. insulin, the mammalian target of rapamycin (mTOR), AMP-activated protein kinase (AMPK) signaling and the insect hormone biosynthesis pathways at 24 h (*Supplementary figure 5f, g, h, i*), steroid hormone biosynthesis, insulin signaling pathways at 48 h (*Supplementary figure 5j, k*), and the mTOR, protein digestion and absorption, and steroid hormone biosynthesis pathways at 72 h. These results may help to explain why HaDV2-positive individuals developed more quickly than non-infected insects.

### HaDV2 decreased the effect of Bt on *H. armigera*

There was a total of 1573 significant differentially expressed genes (DEGs) in *HaDV2-negative* insects after exposure to Cry1Ac (673 up- and 900 down-regulated). We focused on DEGs and pathways related to Bt resistance and immune systems. Seven ABC transporter genes, which are related to immunity, were differentially expressed (4 up- and 3 down-regulated) (*Supplementary figure 6a*); eight trypsin genes, which are related to the conversion of the protoxin to activated toxin, were differently expressed (6 up- and 2 down-regulated) (*Supplementary figure 6b*); two carboxylesterase genes, which are related to the detoxification of Bt by the insect, were up-regulated; and two Bt toxin receptors genes, ALP and APN, which are related to Bt resistance, were down-regulated (*Supplementary figure 6c*). Genes in the mitogen-activated protein kinase (MAPK) signaling pathway, which is related to antimicrobial immune response, was also significantly down-regulated (*Supplementary figure 6d*).

In contrast, there were only 249 significant DEGs in *HaDV2-positive* insects after exposure to Cry1Ac (165 up- and 84 down-regulated). One trypsin gene was down-regulated; genes in the ascorbate and aldarate metabolism pathway, which is related to carbohydrate metabolism were up-regulated; genes in drug metabolism - cytochrome P450 pathway, metabolism of xenobiotics by cytochrome P450 pathway and drug metabolism – and other enzymes pathways, which are related to detoxification, were also up-regulated (*supplementary figure 6e*).

## Discussion

Commercialization of transgenic Bt-crops has brought some significant benefits to farmers (*Carriere et al., 2003; Cattaneo et al., 2006; Lu et al., 2012; Shelton et al., 2002; Wu et al., 2008*). These Bt-plants successfully control target insect pests and have resulted in the reduced use of insecticides in the field (*Bravo et al., 2011; Lu et al., 2012; Wu et al., 2008*). Here, we show that HaDV2 infection enhances resistance to Cry1Ac in both susceptible and resistant strains of *H. armigera.* We also show that there appears to have been strong positive selection for HaDV2-infected *H. armigera* in populations exposed to Cry1Ac-cotton in the field, especially (but not exclusively) when this exposure has been for prolonged periods of time. Intensive monitoring in two provinces where Bt-cotton has been grown over a number of years showed a strong temporal increase in HaDV2 infection levels, from around 40% in 2007 to nearly 100% in 2016, with high levels of HaDV2 infection being associated with higher larval growth rates on Bt-cotton in the field. Indeed, at locations where Bt-cotton is not grown, *H. armigera* show much lower rates of HaDV2-infection (12 - 44% versus 81 - 90%), consistent with weaker selection for HaDV2-infected *H. armigera* in the absence of Bt-crops. Moreover, the prevalence of HaDV2 infection increased with the number of years since Bt-cotton was first introduced to a province and with the proportion of cotton grown that is transgenic, consistent with Bt-cotton being a key selection pressure. There were no significant relationships between HaDV2 prevalence and the proportional cover of any other main crops grown locally (cotton, rice, corn, wheat, beans, tubers, oil-producing crops and vegetables) or with any of the environmental variables tested (rainfall, temperature and altitude). Thus, our data suggest that the Cry1Ac-toxin selected for HaDV2-infected *H. armigera* following exposure to Bt-cotton and our RNA-Seq analysis suggests that the increased resistance of HaDV2-infected insects is caused, in part at least, by the activation of a series of immune pathways and pathways that enhance development.

The emergence and rapid spread of a symbiont through a host population has been observed previously (*Himler et al., 2011; Turelli & Hoffmann, 1991; Weeks et al., 2007*). For example, the bacterial symbiont, *Rickettsia* sp. nr. *bellii*, swept through a population of invasive sweet potato whitefly, *Bemisia tabaci*, in just six years (*Himler et al., 2011*). Although we do not know when HaDV2 first infected *H. armigera* in China, in two areas where Bt-cotton is grown extensively, and where nearly 100% of all insects are currently infected with HaDV2, the prevalence of the densovirus was as low as 40% in 2007, when the earliest samples were collected (*Figure 4*). Moreover, extrapolation of the observed temporal trends (*Figure 4a, b and 5*) would suggest that emergence of HaDV2 may have coincided with the introduction of Bt-cotton in China two decades ago. This apparently rapid spread is perhaps not surprising given the strong fitness benefits of carrying the densovirus (amounting to a ~40% increase in *H. armigera R*_0_) (*Figure 2*).

Rather than being a recent introduction to *H. armigera* in China, it is possible that the association between the HaDV2 and its host is more ancient, but that the increase in HaDV2 prevalence in recent years is because it has only recently evolved from a parasitic or commensal relationship into one that is more mutualistic (*Weeks et al., 2007*), perhaps under direct or indirect selection from Bt-cotton. Indeed, most of the densoviruses that have been studied thus far have tended to be at the parasitic end of the symbiotic spectrum and some of these have even been promoted as potential biological pesticides (*El-Far et al., 2012*). However, the observation that densoviruses tend to be parasitic in nature is likely biased by the fact that viruses with notable pathological effects on their hosts are more likely to be reported (*Roossinck, 2011, 2015; Webster et al., 2015; Xu et al., 2020*). With the advent of modern molecular methods such as suppression subtractive hybridization (SSH) and RNA-seq, it has become apparent that there are many ‘good viruses’ that have failed to be discovered due to their relatively benign, or even beneficial, effects on their hosts (*Roossinck, 2011, 2015; Webster et al., 2015; Xu et al., 2020*), including HaDV2 (*Xu et al., 2014*). Although there are an increasing number of studies suggesting that symbiotic microbes can impact an insect host’s ability to combat pathogens and parasites (*Graham et al., 2012; Hedges et al., 2008; Oliver et al., 2003; Xu et al., 2020*), as far as we are aware, this is the first field study to provide support for the notion that a symbiotic virus could protect a crop pest from Bt toxin.

Regardless of whether or not larvae are exposed to Cry1Ac, HaDV2 appears to enhance the fitness of both Bt-resistant and Bt-susceptible *H. armigera* (*Xu et al., 2014*) (*Figure 2*). Theoretically, the infection rate should naturally increase in the *H. armigera* population both in Bt-cotton and non-Bt-cotton regions because of the mutualistic relationship between HaDV2 and its’ host (*Xu et al., 2014*). We monitored HaDV2 infection rates at 38 locations across sixteen provinces during the period 2014-2016. Indeed, our data showed that HaDV2 infection level increased both in Bt-cotton and non-Bt-cotton areas. However, with the exception of samples from Changde (Hunan province), the other 37 locations showed consistent trends that HaDV2 infection rates were significantly higher in Bt-cotton areas than in non-Bt-cotton areas. Previously, we also quantified the infection rate of HaDV2 in samples collected from Changde in three years (*Xu et al., 2014*). The final infection rate in samples from Changde in the four years was 45%, which is higher than the infection rate of samples from non-Bt-cotton planting regions (26%). Changde is located at the southernmost point of Bt-cotton growing range, where the planted cotton included both non-Bt and Bt cotton and the fluctuation of HaDV2 infection rate in Changde might be due to both the lower pressure of Bt cotton and the migration of *H. armigera* between Bt and non-Bt cotton growing areas. Previously, we showed that there was evidence for a significant decline in HaDV2 prevalence from 2008 to 2012, especially in the migrating moths collected in Yantai (Beihuang Island), which is located in Bohai Sea between Bt and non-Bt-cotton areas and there were no local populations of *H. armigera* (*Feng et al., 2004; Feng et al., 2005; Feng et al., 2009; Xu et al., 2014*). If we exclude the data from Yantai, however, there was no obvious temporal trend in HaDV2 prevalence between 2008 and 2012 (85%, 77%, 83%, 73% and 75% respectively). Other factors might also result in variation in HaDV2 infection rates. For example, there are four generations of *H. armigera* per year in northern China and the fourth generation overwinters as diapausing pupae, and if *H. armigera* grows too fast due to HaDV2 infection, it will overwinter at an inappropriate state which may lead to higher mortality of HaDV2-positive individuals (*Mu et al., 1995; Zhang & Li, 2001*). The climate and crop structures of the locations for our samples collected in this study have been relatively constant over the last few decades and data analyses supported the notion that HaDV2 prevalence was not correlated with environmental variables or the areas of any other crops being grown. These results support the notion that Bt-cotton had played an important role in the increasing HaDV2 infection rate in field populations of *H. armigera*, and suggest the possibility that HaDV2 may threaten the future control of *H. armigera* by Bt.

We found some evidence to support this notion from LC50-bioassays using synthetic diets containing Cry1Ac protoxin, with the benefits of hosting HaDV2 being greatest for the most Bt-resistant strains (*Figure 1*), suggesting synergistic effects of Bt-resistance and HaDV2-infection. However, laboratory trials using non-Bt and Bt-cotton plants suggested that the relative fitness benefits of harboring HaDV2 were independent of exposure to Bt-cotton and of Bt-resistance levels (at least in terms of larval growth rates) (*Supplementary table 6*). This is at odds with the observation that HaDV2 infection levels in the field are higher when populations are exposed to Bt-cotton for prolonged periods, and may indicate that LC50-bioassays are a more reliable measure of relative fitness than larval growth on Bt-cotton plants in the laboratory. Alternatively, it is possible that there are added benefits to HaDV2-infection for *H. armigera* exposed to Bt-cotton in the field that our laboratory bioassays cannot fully capture, such as enhanced resistance to predators and parasitoids, or reduced exposure to chemical pesticides. In this latter regard, there may be parallels with the increase in outbreaks of mirid bugs (Heteroptera: Miridae) following wide-scale adoption of Bt-cotton in China and the parallel reduction in pesticide application rate on crops (*Lu et al., 2010; Zhang et al., 2018*).

The evolution of resistance to Bt-toxins could endanger the utility of this GM technology (*Gahan et al., 2001, 2007and 2010; Heckel, 2012; Tabashnik, 2015; Tabashnik et al., 2008 and 2013*). However, as observed here, resistance to Cry toxins is often associated with significant fitness costs that are likely to delay the establishment of Bt-resistant populations under field conditions (*Gassmann et al., 2009; Horikoshi et al., 2016; Liu et al., 2017; Raymond et al., 2005; Santos-Amaya et al., 2017*). Interestingly, we show here that Bt-resistant strains with different resistance mechanisms, such as mutations affecting ABCC2 (strain LF60) or cadherin (strain 96CAD) (*Liang et al., 2008; Xiao et al., 2014*), incurred significantly reduced fitness costs when they were infected with HaDV2. The reduction of fitness costs by HaDV2 infection could speed up the establishment of resistant populations. Thus, a potential implication of these results is that the increased prevalence of HaDV2 infections under field conditions could impact the evolution of *H. armigera* populations resistant to Bt-cotton.

The temporal increase in HaDV2-infection levels observed in Xiajin and Anci, two cotton-growing counties >300 km apart in north-east China, was associated with parallel temporal increases in relative growth rates (RADR; Fig. 4c, d). In a previous study of these two populations, the temporal increase in RADR between 2002 and 2014 was ascribed to evolved genetic resistance to Cry1Ac in *H. armigera*, despite the failure to identify the key genetic locus conferring resistance in these populations (An et al., 2015). The relatively strong correlation between the annual prevalence of HaDV2 in *H. armigera* and their mean annual RADR in each of these populations (*Figure 4e,f*) suggests that the putative increase in genetic resistance over this period may, at least in part, be due to the phenotypic expression of HaDV2-infection. Future studies will seek to tease apart these genetic and phenotypic effects. The fact that the densovirus is mostly located in the insect’s fat body suggests that it may play a role in *H. armigera* development (*Xu et al., 2014*). Lepidopteran larvae tend to become more resistant to pathogens and toxins as they age and grow, a phenomenon known as developmental resistance (*Engelhard & Volkman, 1995*). HaDV2-infected *H. armigera* not only accumulate more fat body than non-infected individuals, but they also grow faster and larger (*Xu et al., 2014*). Therefore, developmental resistance could provide a potential mechanism to explain the enhanced resistance of densovirus-infected insects. HaDV2-infected larvae have more fat body than non-infected insects (*Xu et al., 2014*) and the fat body plays an important role in insect immunity as it is a major site for the production and secretion of antimicrobial peptides (*Hoffmann, 2003; Lemaitre & Hoffmann, 2007*), so immunological resistance is another potential mechanism by which HaDV2 may enhance resistance to both baculovirus and Cry1Ac.

To explore further the molecular interaction between the HaDV2 and its host, we performed an RNA-seq experiment using *H. armigera* larvae. Due to the observed effects of HaDV2 on its host, we focused on molecular pathways related to immunity and development (*Jindra et al., 2013; Kingsolver & Hardy, 2012; Kingsolver et al., 2013; Lin & Smagghe, 2018; Shields, 2017*). In *H. armigera* larvae at 24 and 48 h, the Jak-STAT pathway was significantly enriched by up-regulation in HaDV2-positive individuals. In fruit flies and mosquitoes, this pathway controls the expression of genes in response to infection with a range of viruses and bacteria including Drosophila C virus, dengue virus, West Nile virus, *Escherichia coli* and *Micrococcus luteus* (*Barillas-Mury et al., 1999; Marques & Imler, 2016*). The Jak-STAT pathway was not significantly enriched in larvae at 72 h, however, pathways related to other antimicrobials were, e.g. the lysosome and MAPK signaling pathways (*Takano, et al., 2019; Watts, 2012*). Previously, we found that HaDV2-positive individuals developed faster than non-infected insects (*Xu et al., 2014*). Interestingly, the transcriptome data suggest that several pathways related to development are also significantly enriched, including the insect hormone synthesis, insulin, mTOR and AMPK signaling pathways (*Bland et al., 2010; Jindra et al., 2013; Lin & Smagghe, 2018; Saxton & Sabatini, 2017*). We did not observe the differential expression of genes playing pivotal roles in Bt resistance of *H. armigera*, e.g. cadherin, aminopeptidase N (APN) and alkaline phosphatase (ALP) (*Chen et al., 2015; Gahan et al., 2001; Zhang et al., 2009*), suggesting that HaDV2 may not enhance resistance to Bt using traditional molecular resistance pathways, though we did observe that the ABC transporter pathways, which has been reported to be associated with Bt-resistance (*Bretschneider et al., 2016; Dermauw & Van Leeuwen, 2014*), were significantly up-regulated at 48 h post challenge.

Generally, the mode of action of Bt toxins agree that Bt protoxins are first converted to activated toxins by midgut proteases, then the toxins bind to the midgut receptors, e.g. CAD, APN and ALP, finally leading to insect death (*Xiao & Wu, 2019*). A change in any of these steps will lead to insect resistance, such as mutations in genes for toxin activation, toxin-binding, and changes in insect immunity (*Xiao et al., 2014; Zhang et al., 2009*). To understand the responses of *H. armigera* to Bt toxins with or without HaDV2, we conducted RNA-seq after larvae had been exposed to Cry1Ac. In insects lacking HaDV2, Cry1Ac increased the activation of Bt protoxins by increasing the expression level of proteases, such as trypsin. In a defensive response to Cry1Ac, receptor genes of *H. armigera*, such as APN and ALP, were significantly down-regulated to reduce the toxin-binding capacity. Carboxylesterase genes and most of ABC transporters genes were significantly up-regulated to improve insect immunity, while other pathways related to antimicrobials, e.g. MAPK signaling pathways, were significantly down-regulated, reducing resistance to Bt toxin (*Yang et al., 2020*). In contrast, when *H. armigera* larvae were infected with HaDV2, these genes and pathways were generally not significantly enriched after Cry1Ac exposure: one trypsin gene was down-regulated and some genes involved in drug metabolism were up-regulated to increase the metabolism of xenobiotics, improving Bt resistance. Through RNA-seq analysis, we did not find significant up-regulation or down-regulation of genes known to be related to typical Bt resistance. Thus, it is difficult to perform further experiments, e.g. CRISPR. In spite of this, our data still support the notion that HaDV2 increased the Bt resistance level of *H. armigera* via related immune mechanisms.

In summary, here we report the rapid increase in the prevalence of a mutualistic virus in a major crop pest and show that its spread is greatest for field populations exposed to Bt-cotton for prolonged periods. Our findings support a novel mechanism by which insects may cope with changes in engineered host plant defences, namely via infection with a symbiotic virus that enhances their resistance to the toxins expressed by Bt-cotton crops. Our study has significant implications for understanding the co-evolution of host-pathogen-symbiont interactions.

## Materials and methods

### Bt toxins

Cry1Ac was obtained as a gift from Dow AgroSciences (Indianapolis, IN) in product formulation MVPII (20%). To avoid degradation, Cry1Ac protoxin was stored at −80℃. To exclude the possibility of a decline in protoxin potency over time, the susceptible strains (LF) were used as internal control in different years (*Cao et al., 2014*).

### Laboratory strains

Different *H. armigera* strains were used: two susceptible strains (LF and 96S) and eight Cry1Ac-resistant strains (BtR, 96CAD, LFC2, LF5, LF30, LF60, LF120, LF240), with different resistance ratios due to different resistant mechanisms (*Liang et al., 2008; Xiao et al., 2014*). The Bt-susceptible strains, LF and 96S, were collected from Langfang, Hebei Province, in 1998, and Xinxiang, Henan Province, in 1996, respectively. They had been continuously cultured in the laboratory without exposure to Bt toxin. The resistant strains, were selected from the susceptible strain on artificial diets (*Cao et al., 2014; Liang et al., 2008*); 96CAD (with a cadherin mutation) and LFC2 (with an ABCC2 mutation, unpublished) are two near-isogenic lines isolated from BtR and LF60 resistant strains, respectively (*Xiao et al., 2017*). All strains were reared on artificial diet. Rearing, selection, and bioassays were conducted at 25±1°C, photoperiod 14L:10D, and 75±10% relative humidity.

### Collection of field strains

Adult female bollworm moths were collected from a 1000 W light traps in two counties 360 km apart: Xiajin County, Shandong Province (with an intensive cotton planting area), and in Anci County, Hebei Province (with a multiple-crop farming system that includes cotton). The history of Bt-cotton deployment in the two locations during 2007 – 2013 has been reported previously (*An et al., 2015*). In the two counties*, H. armigera* female moths were trapped from June to October in all years, as previously reported (*An et al., 2015*).

### HaDV2 preparation

HaDV2-containing liquid fluids was prepared as described previously (*Xu et al., 2014*). Briefly, the collected individual moths were ground in liquid nitrogen completely. Ten micrograms of powder were transferred to a clean 1.5 mL tube and DNA was extracted. PCR reactions were undertaken to detect the presence of HaDV2 using specific primers, HaDV-F: GGATTGGCCTGGGAAATGAC and HaDV-R: CGTTGTTTTTATATCCGAGG. The remaining debris of HaDV2-positive individuals was transferred to 1 ml PBS buffer on ice. The homogenate was centrifuged and the liquid supernatant was subsequently filtered with 0.22 μm membrane filter (Sigma). The HaDV2-containing filtered liquid (200 μL per tube) was collected and stored immediately at −80 °C. Quantification of the virus was performed using the qPCR method (*Xu et al., 2014*).

### Detection of HaDV2 in wild populations of *H. armigera*

Samples of larvae or adults were collected at 12 locations in 2014, 20 locations in 2015 and 18 in 2016 (*Supplementary table 7*). Infection rates with HaDV-2 were determined using the PCR method described in Xu et al. (*2014*).

### Bioassays

*H. armigera* strains were reared and assessed simultaneously for susceptibility to Cry1Ac, as previously reported (*Liang et al., 2008*). In brief, the susceptibility was evaluated by feeding *H. armigera* larvae with artificial diet containing different concentrations of Cry1Ac toxin. Newly-hatched larvae were first orally inoculated with either filtered-liquid containing HaDV2, or filtered-liquid from uninfected individuals. After that, they were placed in each treatment Petri dish for 2 days to ensure that larvae ingested the treated diet. They were then transferred individually on Cry1Ac-contaminated artificial diet in plastic wells (depth: 1.5 cm, vol. 3 ml) in 24-well plates. The plates were covered with a plastic lid to prevent escape and 72 larvae were tested per Cry1Ac concentration. After seven days, the mortality was recorded and body mass of individuals that were still alive was measured. Both dead larvae and those with a body mass of less than 5 mg were recorded as dead. For each strain, the median lethal concentration (LC_50_) value was determined using a Probit analysis (POLO Plus LeOra Software, Berkeley) and the resistance level to Cry1Ac indicated by the resistance ratio was calculated with LC_50_ of the tested strains divided by the LC_50_ of the susceptible strain.

### Assessment of fitness cost

To test the impact of HaDV2 infection on the fitness cost of its host, life table parameters of the different strains were analyzed on artificial diet. The oral inoculation method was the same as previously described. Thirteen fitness components (life history parameters) were obtained with life-table techniques (*Supplementary table 3 and 4*). The components considered were: (i) survival rate from the 1^st^ to 5^th^ larval instar (0 – 1), (ii) survival rate from the 5^th^ instar to the pupal stage (0 –1), (iii) duration of larval development (days), (iv) pupal weight (mg), (v) female pupal development period (days), (vi) male pupal development period (days), (vii) adult emergence rate (0 – 1), (viii) proportion of females (or sex ratio) (0 – 1), (ix) copulation rate (0 – 1), (x) female adult longevity (days), (xi) male adult longevity (days), (xii) fecundity (number of eggs laid) and (xiii) hatching rate (0 – 1). The net reproductive rate (*R*_0_) was calculated for each *H. armigera* strain (*Cao et al., 2014*) as *R*_0_ = N_t+1_/N_t_, where N_t_ is the population size of the parent generation and N_t+1_ is that of its next generation. When *R*_0_> 1, it indicates a higher number of female offspring produced than that of the parental females. We used thirty larvae per repeat and three repeats for each treatments.

To analyze the fitness cost associated with Cry1Ac-resistance, the larval weight of the different strains was analyzed on Bt/non-Bt-cotton. Newly-hatched larvae were first orally inoculated with either filtered-liquid containing HaDV2, or filtered-liquid from non-infected individuals. After that, they were placed in each treatment Petri dish for 2 days to ensure that larvae ingested the treated diet. They were then transferred individually onto Bt-cotton (Zhongmian 29) or non Bt-cotton (Zhongmian 24) leaf in plastic wells in 24-well plates. After nine days, the weight of individuals was measured. Each treatment had 48 larvae with 3 repeats.

### Bioassay of F1 generation on Bt and non-Bt diets

To obtain F1 generation offspring of each of the laboratory strains, one virgin adult male and one virgin adult female were paired in a 250 mL clear plastic cup. Adult females collected from the field were placed individually into 250 mL clear plastic cups and covered with gauze to provide a substrate for egg laying. Eggs were collected on a daily basis. At larval hatch, 24 neonates from each female line were placed on non-Bt diet, and 24 neonates were placed on Cry1Ac-containing diet (1 μg of Cry1Ac/g diet). The concentration of Bt Cry1Ac protoxin was 1.0 μg/g diet. The composition of the diet was previously described (*Chen et al., 2015*). Rearing, selection and bioassays were conducted at 27 ± 2 °C with a photoperiod of 14:10 h L:D and a relative humidity of 75 ± 10%. Larvae were scored for developmental stage after 6 days. Larval instar was determined on the basis of head capsule and body size.

### Analyzing the effects of the HaDV2 on its hosts by transcriptome analysis

To determine the effect of HaDV2 on *H. armigera* at a transcriptomic level, we collected samples of the HaDV2-negative and -positive individuals from single pairs of *H. armigera* and performed RNA-seq, using larvae at 24 h, 48 h and 72 h after hatching. Briefly, newly hatching larvae were fed on filtered liquid with/without HaDV2 as described previously (*Xu et al., 2014*). Samples were collected at 24 h, 48 h and 72 h and there were two groups of 30 individuals for each group per trial (*Supplementary table 8*). The Trinity (v2.0.6) (*Grabherr et al., 2011*) software were used to assemble the clean reads with default parameters. BLASTx was performed to align the assembled contigs from Trinity to the database of NR, String, Swissprot and KEGG for functional annotation. The e-value cut-off was set at 1E-5 for further analysis. For gene expression analysis, the reads from 12 libraries were mapped to the assembled contigs using Bowtie 0.12.7 (*Langmead et al., 2009*) with no more than 2 mismatches within the first 28 bp. The read counts accumulated on the contigs were normalized as fragments per kilobase of exon model per million mappedreads (FPKM) values (*Trapnell et al., 2010*). Quantitative analysis for each unigene was estimated using FPKM values by RSEM (v1.1.17) software (*Li & Dewey, 2011*) with default parameters. Then the R package DEseq2 (*Love et al., 2014*) was used to get the significantly differential expressed unigenes at different comparisons. The threshold to determine the significantly differential expressed unigenes was ‘fold change ≥ 1.5 and the P < 0.05’. The hierarchical clustering method was applied to analyze the expression pattern of significantly differentially expressed unigenes in different samples. Significantly enriched KEGG pathways were identified using the Fishers exact test (p-value<0.05) (*Klopfenstein et al., 2018*).

### RNA sequence analysis

Offspring from a single uninfected breeding pair were reared to produce the N-strain (uninfected) laboratory culture. Neonate N-strain larvae were first orally-inoculated with either filtered-liquid containing HaDV2 (10^8^ copies/μl), or filtered-liquid from non-infected individuals (control). One hundred N-strain neonates were placed in each treatment Petri-dish for 2 days to ensure that larvae ingested the treated diet. They were then transferred to a 24-well plate (one individual per well: diameter = 1.5 cm; height = 2 cm) containing the artificial diet containing 1 μg/ml Cry1Ac toxins. After 48 hours, the larvae were collected and stored at −80 °C for transcriptome sequencing. Twenty larvae were pooled together as a sample. A total of 12 samples were sequenced including three replications of four treatments (HaDV2-Bt-, HaDV2-Bt+, HaDV2+Bt-, HaDV2+Bt+).

### Quantification and Statistical Analysis

The relative average development rates, RADR, were calculated according to An et al. (*2015*). In brief, the RADR for a line was calculated as the average body length rating of larvae from that line reared on the Bt diet divided by the average rating of larvae from that line reared on the non-Bt diet. Multivariate 2 factor variance analysis was conducted with IBM SPSS 20. Student’s t-test or ANOVA with Tukey-test *post hoc* comparisons were used to determine the level of significance. Simple linear regression model was used to analyze the temporal trends of *H. armigera,* including RADR and HaDV2 infection rate, from 2007 to 2016. Simple linear regression model was also used to analyze the association relationship of RADR and HaDV2 infection rate in different years. Logistic regression, using the R statistical package (version 3.3.3) (*R Core Team, 2017*), was used to test the association between HaDV2 infection rate and environmental variables (rainfall, temperature and altitude the proportional cover of the main crops (cotton, rice, corn, wheat, beans, tubers, oil-producing crops and vegetables, the Bt status of the cotton crop, and the number of years since the introduction of Bt-cotton to a province (defined as the year in which Bt-cotton comprised at least 10% of the cotton grown in a province).

## Acknowledgments

We thank Bruce E. Tabashnik (Department of Entomology, University of Arizona, USA) and Ling Wang (Institute of Plant Protection and Soil Fertility, Hubei Academy of Agricultural Sciences, China) for their comments on data analysis.

## Declaration of Interests

M.S. and A.B. are coauthors of a patent on modified Bt toxins, “Suppression of Resistance in Insects to *Bacillus thuringiensis* Cry Toxins, Using Toxins that do not Require the Cadherin Receptor” (patent numbers: CA2690188A1, CN101730712A, EP2184293A2, EP2184293A4, EP2184293B1, WO2008150150A2, WO2008150150A3). Authors declare that no competing interests exist.

## Data availability statement

All raw sequence data generated during this study have been deposited at NCBI as BioProject under accession PRJNA638220 and PRJNA638972.

## Supplementary Information

**Supplementary table 1:**
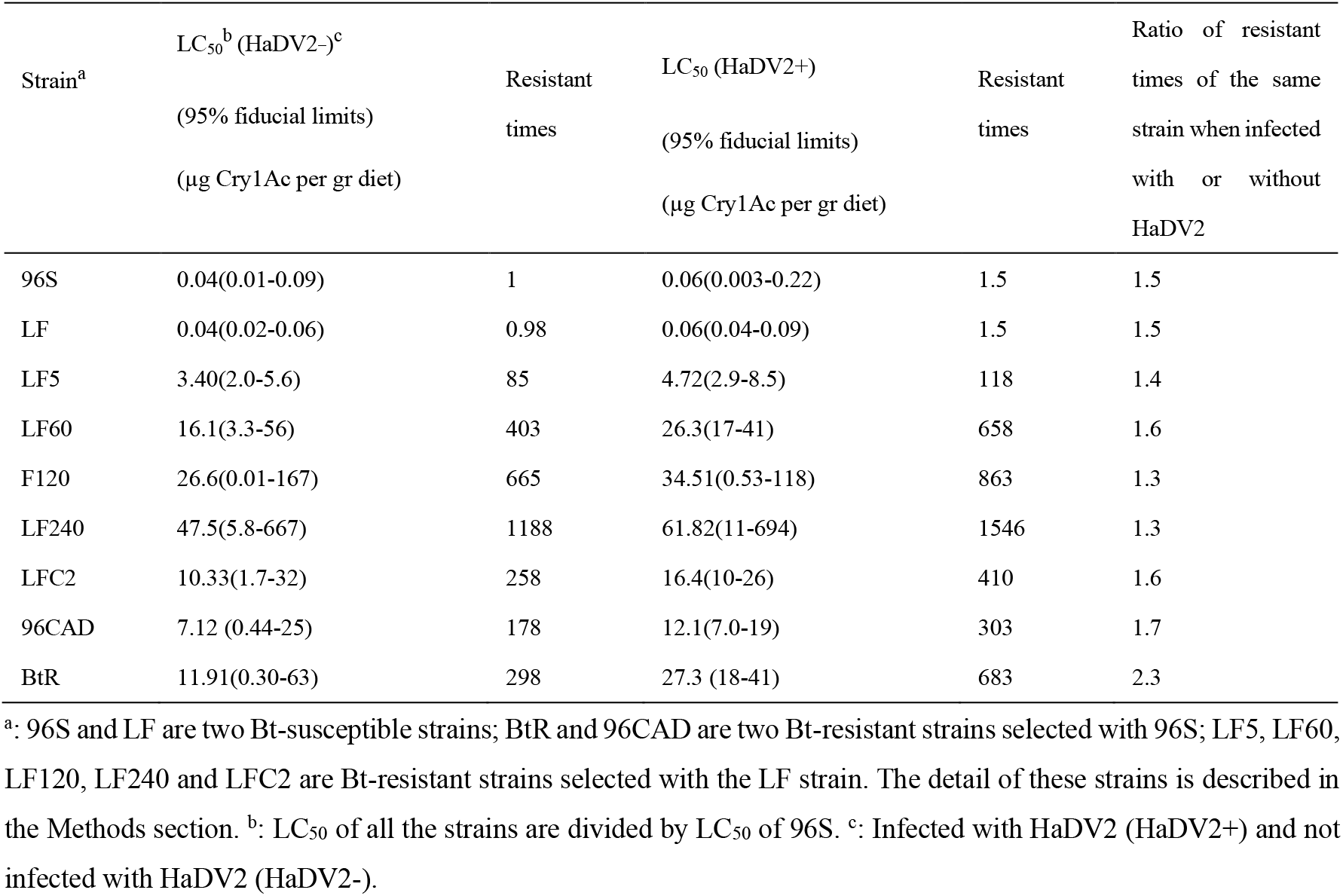
Bt toxin sensitivity test of different *H. armigera* strains with or without HaDV2 infection.

**Supplementary table 2:**
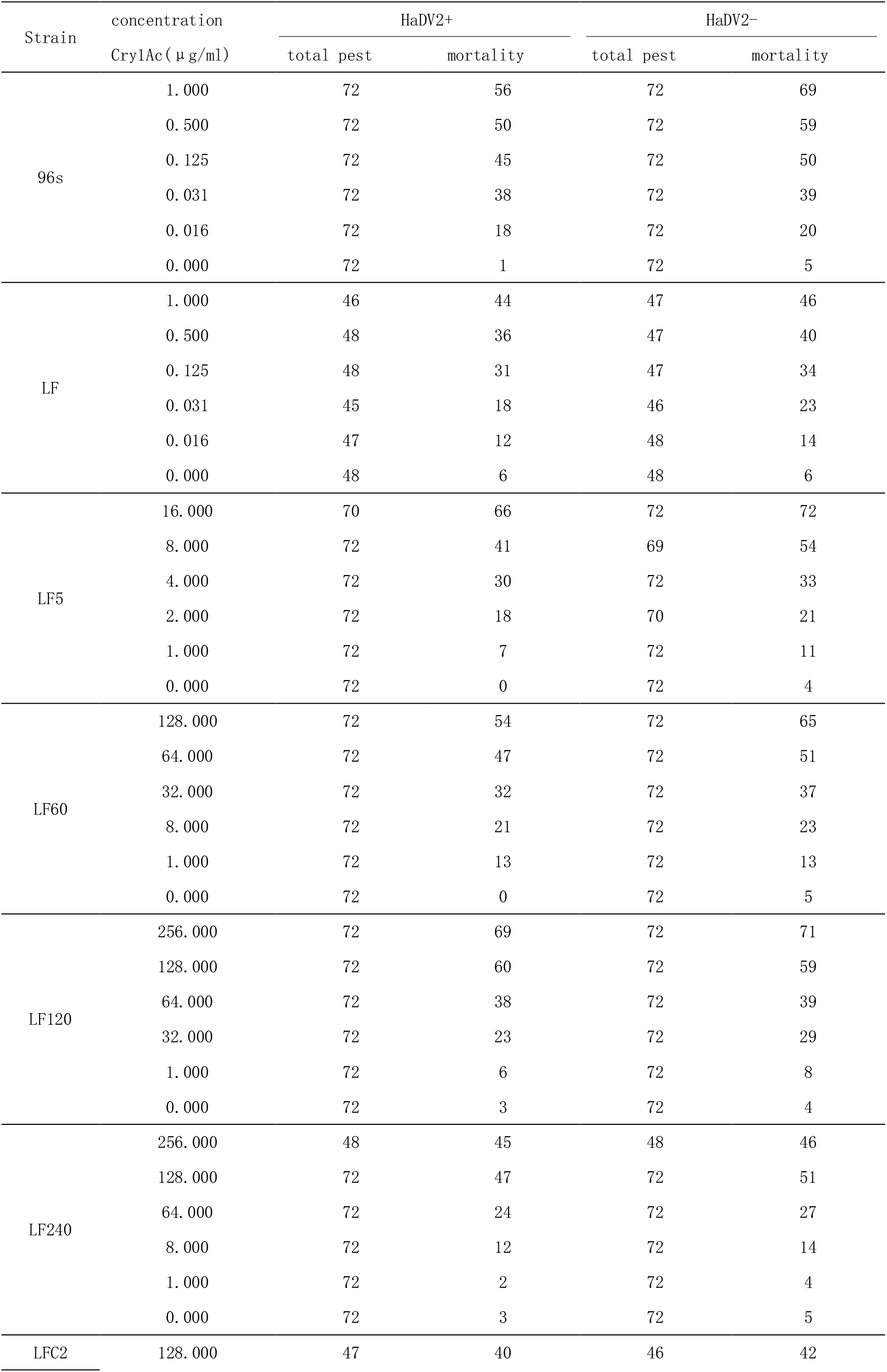
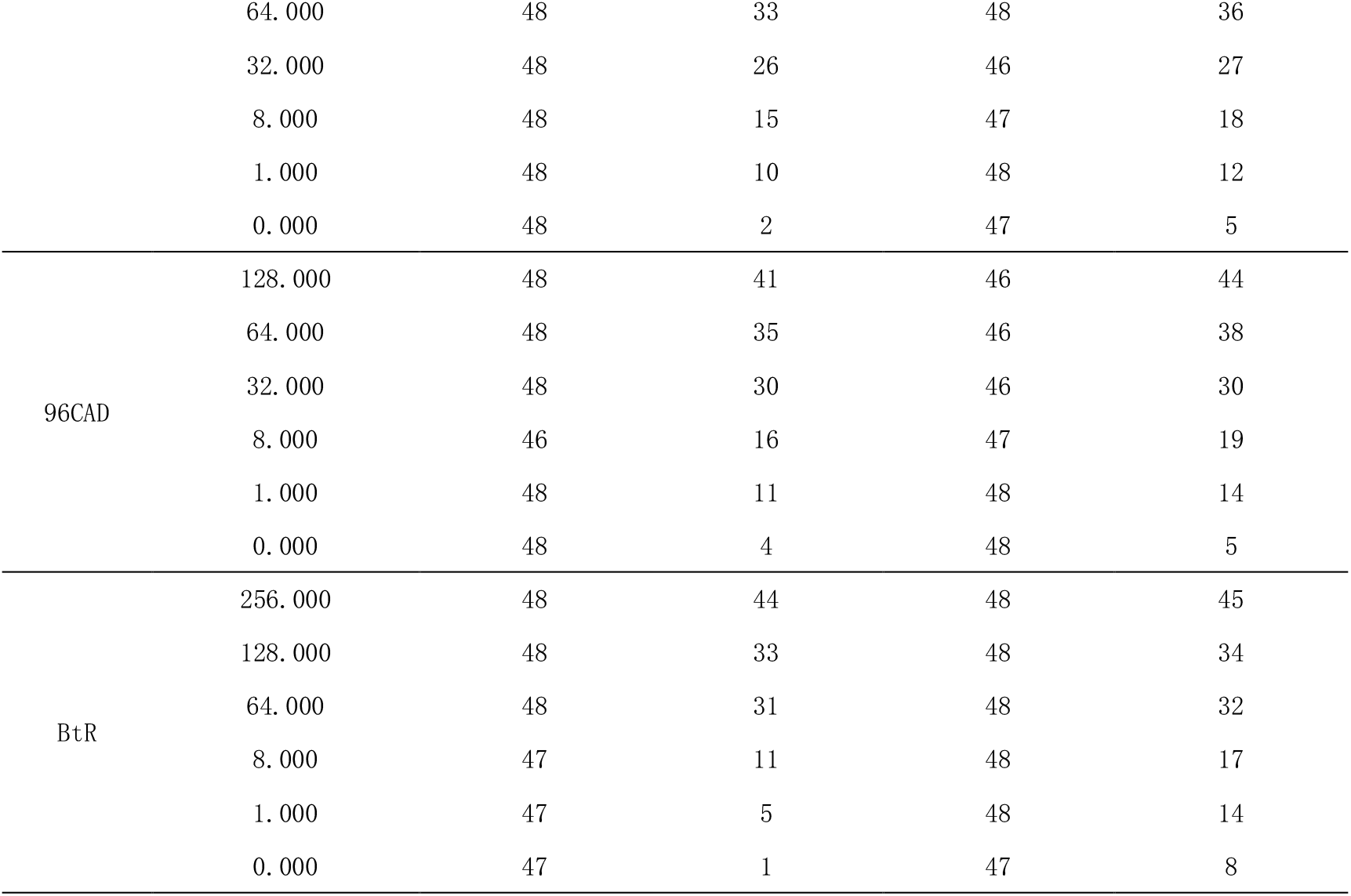
Mortality changes with Bt toxin concentration test of different *H. armigera* strains with or without HaDV2 infection.

**Supplementary table 3.** Comparing of the effects of HaDV2 on fitness components of LF, LF5, LF60 and LF240. LF is susceptible strain; LF5, LF60 and LF240 are Bt resistant strains selected with LF strain. Significant differences (ANOVA followed by Tukey’s HSD test) between each strain with or without HaDV2 infestation are indicated by different letters (p < 0.05). Insects were reared on artificial non Bt-diet. D+ stand for infected by HaDV2, D- stand for non-infected by HaDV2. (1-5) means the survival rate from the first star to the 5th star; 5-p: from the 5th star to pupa; Proportion FA: the rate of female divided male.

| *H. armiger a* strains | Survival rate (1-5) | Survival rate (5-p) | Larval period (days) | Pupal weight (mg) | Female pupa period (days) | Male pupa period (days) | Emergence rate | Proportion FA | Longevity of female adult | Longevity of male adult | Number of eggs produced per female | Hatchling rate |
| --- | --- | --- | --- | --- | --- | --- | --- | --- | --- | --- | --- | --- |
| LF D+ | 0.92±0.02 a | 0.94±0.03 a | 16.76±0.24 f | 236.11±9.55 a | 11.23±0.17 e | 13.03±0.36 e | 0.92±0.032 a | 0.51±0.04 a | 8.07±0.33 a | 9.52±0.27 abc | 477.29±10.46 a | 0.69±0.02 a |
| LF D- | 0.89±0.02 abc | 0.91±0.03 abc | 17.43±0.19 e | 224.68±21.61 ab | 11.94±0.22 cd | 13.78±0.31 c | 0.92±0.024 a | 0.51±0.05 a | 7.61±0.34 a | 9.92±0.40 ab | 402.17±11.51 b | 0.62±0.02 bc |
| LF5 D+ | 0.90±0.02 ab | 0.94±0.03 a | 17.38±0.14 e | 223.27±8.02 ab | 11.58±0.24 de | 13.29±0.24 de | 0.92±0.017 a | 0.49±0.02 a | 8.10±0.25 a | 10.33±0.32 a | 408.61±17.23 b | 0.64±0.02 ab |
| LF5 D- | 0.87±0.02 abcd | 0.92±0.02 bc | 17.92±0.11 cd | 215.25±5.70 b | 12.08±0.22 cd | 13.63±0.25 cd | 0.91±0.024 a | 0.50±0.02 a | 7.74±0.27 a | 9.45±0.34 abc | 348.77±12.81 c | 0.59±0.02 bc |
| LF60 D+ | 0.85±0.02 abcd | 0.85±0.03 abc | 17.74±0.14 de | 222.45±6.33 ab | 11.73±0.21 de | 13.60±0.33 cd | 0.91±0.031 a | 0.50±0.02 a | 7.79±0.32 a | 9.14±0.26 bc | 408.32±18.51 b | 0.63±0.02 bc |
| LF60 D- | 0.81±0.03 cd | 0.85±0.03 bc | 18.20±0.11 c | 224.34±6.42 ab | 12.41±0.20 bc | 14.28±0.29 b | 0.91±0.023 a | 0.49±0.03 a | 7.63±0.33 a | 8.95±0.30 c | 333.45±11.98 c | 0.58±0.02 c |
| LF240 D+ | 0.83±0.03 bcd | 0.83±0.04 c | 19.03±0.15 b | 151.13±4.15 bc | 12.93±0.15 ab | 14.23±0.27 b | 0.85±0.029 b | 0.49±0.02 a | 7.83±0.25 a | 8.76±0.31 c | 340.12±12.30 c | 0.50±0.02 d |
| LF240 D- | 0.80±0.05 d | 0.83±0.04 c | 19.61±0.17 a | 126.25±4.38 c | 13.31±0.14 a | 14.68±0.25 a | 0.80±0.045 b | 0.51±0.03 a | 7.43±0.21 a | 8.57±0.31 c | 292.30±9.52 c | 0.42±0.02 e |

**Supplementary table 4.** Analysis of variance for fitness parameters of 4 cotton bollworm strains (LF, LF5, LF60, and LF240)

| Source | Dependent variable | Type III sum of squares | df | Mean square | F | P |
| --- | --- | --- | --- | --- | --- | --- |
| Model | Survival rate (1-5) | 0.118 | 12 | 0.010 | 2.336 | 0.025 |
|  | Survival rate (5-p) | 0.157 | 12 | 0.013 | 2.977 | 0.006 |
|  | Larval period (days) | 7853.346 | 25 | 314.134 | 1.233 | 0.205 |
|  | Pupal weight | 28274.921 | 273 | 103.571 | 0.999 | 0.607 |
|  | Female pupa period (days) | 2235.348 | 24 | 93.139 | 0.714 | 0.836 |
|  | Male pupa period (days) | 2241.381 | 21 | 106.732 | 0.868 | 0.633 |
|  | Emergence rate | 0.203 | 12 | 0.017 | 6.835 | 0.000 |
|  | Proportion FA | 0.019 | 12 | 0.002 | 0.285 | 0.988 |
|  | Longevity of female adult | 187.565 | 48 | 3.908 | 1.218 | 0.170 |
|  | Longevity of male adult | 307.072 | 50 | 6.141 | 1.615 | 0.009 |
|  | Number of eggs produced per female | 1148918.561 | 44 | 26111.785 | 4.302 | 0.000 |
|  | Hatching rate | 1.889 | 39 | 0.048 | 4.840 | 0.000 |
| Intercept | Survival rate (1-5) | 35.477 | 1 | 35.477 | 8455.099 | 0.000 |
|  | Survival rate (5-p) | 37.375 | 1 | 37.375 | 8519.698 | 0.000 |
|  | Larval period (days) | 55338.987 | 1 | 55338.987 | 217.264 | 0.000 |
|  | Pupal weight | 86529.972 | 1 | 86529.972 | 834.694 | 0.000 |
|  | Female pupa period (days) | 28229.588 | 1 | 28229.588 | 216.492 | 0.000 |
|  | Male pupa period (days) | 22789.725 | 1 | 22789.725 | 185.415 | 0.000 |
|  | Emergence rate | 38.149 | 1 | 38.149 | 15381.938 | 0.000 |
|  | Proportion FA | 12.070 | 1 | 12.070 | 2145.183 | 0.000 |
|  | Longevity of female adult | 15159.147 | 1 | 15159.147 | 4724.650 | 0.000 |
|  | Longevity of male adult | 23454.485 | 1 | 23454.485 | 6169.256 | 0.000 |
|  | Number of eggs produced per female | 33712416.161 | 1 | 33712416.161 | 5554.433 | 0.000 |
|  | Hatching rate | 74.448 | 1 | 74.448 | 7440.300 | 0.000 |
| RL | Survival rate (1-5) | 0.074 | 3 | 0.025 | 5.850 | 0.002 |
|  | Survival rate (5-p) | 0.091 | 3 | 0.030 | 6.913 | 0.001 |
|  | Larval period (days) | 2216.175 | 9 | 246.242 | 0.967 | 0.467 |
|  | Pupal weight | 27873.003 | 269 | 103.617 | 1.000 | 0.607 |
|  | Female pupa period (days) | 870.277 | 6 | 145.046 | 1.112 | 0.355 |
|  | Male pupa period (days) | 484.192 | 6 | 80.699 | 0.657 | 0.685 |
|  | Emergence rate | 0.079 | 3 | 0.026 | 10.597 | 0.000 |
|  | Proportion FA | 0.002 | 3 | 0.001 | 0.105 | 0.957 |
|  | Longevity of female adult | 2.135 | 3 | 0.712 | 0.222 | 0.881 |
|  | Longevity of male adult | 76.201 | 3 | 25.400 | 6.681 | 0.000 |
|  | Number of eggs produced per female | 548157.882 | 3 | 182719.29<br>4 | 30.105 | 0.000 |
|  | Hatching rate | 1.159 | 3 | 0.386 | 38.595 | 0.000 |
| HaDV2 | Survival rate (1-5) | 0.011 | 1 | 0.011 | 2.729 | 0.108 |
|  | Survival rate (5-p) | 0.002 | 1 | 0.002 | 0.473 | 0.496 |
|  | Larval period (days) | 1569.766 | 3 | 523.255 | 2.054 | 0.106 |
|  | Pupal weight | 0.000 | 0 | 0 | 0 | 0 |
|  | Female pupa period (days) | 175.156 | 3 | 58.385 | 0.448 | 0.719 |
|  | Male pupa period (days) | 552.641 | 3 | 184.214 | 1.499 | 0.215 |
|  | Emergence rate | 0.003 | 1 | 0.003 | 1.016 | 0.320 |
|  | Proportion FA | 2.083E-6 | 1 | 2.083E-6 | 0.000 | 0.985 |
|  | Longevity of female adult | 3.424 | 1 | 3.424 | 1.067 | 0.303 |
|  | Longevity of male adult | 3.105 | 1 | 3.105 | 0.817 | 0.367 |
|  | Number of eggs produced per female | 291274.783 | 1 | 291274.78<br>3 | 47.990 | 0.000 |
|  | Hatching rate | 0.243 | 1 | 0.243 | 24.301 | 0.000 |
| RL*HaDV2 | Survival rate (1-5) | 0.000 | 3 | 5.092E-5 | 0.012 | 0.998 |
|  | Survival rate (5-p) | 0.001 | 3 | 0.000 | 0.072 | 0.975 |
|  | Larval period (days) | 2494.480 | 12 | 207.873 | 0.816 | 0.634 |
|  | Pupal weight | 0.000 | 0 | 0 | 0 | 0 |
|  | Female pupa period (days) | 1189.612 | 14 | 84.972 | 0.652 | 0.820 |
|  | Male pupa period (days) | 1336.922 | 11 | 121.538 | 0.989 | 0.457 |
|  | Emergence rate | 0.004 | 3 | 0.001 | 0.584 | 0.630 |
|  | Proportion FA | 0.002 | 3 | 0.001 | 0.095 | 0.962 |
|  | Longevity of female adult | 1.287 | 3 | 0.429 | 0.134 | 0.940 |
|  | Longevity of male adult | 19.051 | 3 | 6.350 | 1.670 | 0.174 |
|  | Number of eggs produced per female | 17392.772 | 3 | 5797.591 | 0.955 | 0.415 |
|  | Hatching rate | 0.008 | 3 | 0.003 | 0.282 | 0.839 |
| Error | Survival rate (1-5) | 0.147 | 35 | 0.004 |  |  |
|  | Survival rate (5-p) | 0.154 | 35 | 0.004 |  |  |
|  | Larval period (days) | 101628.692 | 399 | 254.709 |  |  |
|  | Pupal weight | 311.000 | 3 | 103.667 |  |  |
|  | Female pupa period (days) | 36901.857 | 283 | 130.395 |  |  |
|  | Male pupa period (days) | 34415.271 | 280 | 122.912 |  |  |
|  | Emergence rate | 0.087 | 35 | 0.002 |  |  |
|  | Proportion FA | 0.197 | 35 | 0.006 |  |  |
|  | Longevity of female adult | 808.548 | 252 | 3.209 |  |  |
|  | Longevity of male adult | 1037.901 | 273 | 3.802 |  |  |
|  | Number of eggs produced per female | 1414184.518 | 233 | 6069.461 |  |  |
|  | Hatching rate | 2.041 | 204 | 0.010 |  |  |
| Total | Survival rate (1-5) | 35.741 | 48 |  |  |  |
|  | Survival rate (5-p) | 37.685 | 48 |  |  |  |
|  | Larval period (days) | 430366.000 | 425 |  |  |  |
|  | Pupal weight | 117472.000 | 277 |  |  |  |
|  | Female pupa period (days) | 160385.000 | 308 |  |  |  |
|  | Male pupa period (days) | 150869.000 | 302 |  |  |  |
|  | Emergence rate | 38.439 | 48 |  |  |  |
|  | Proportion FA | 12.286 | 48 |  |  |  |
|  | Longevity of female adult | 18955.000 | 301 |  |  |  |
|  | Longevity of male adult | 29513.000 | 324 |  |  |  |
|  | Number of eggs produced per female | 42272760.000 | 278 |  |  |  |
|  | Hatching rate | 88.383 | 244 |  |  |  |
| Corrected total | Survival rate (1-5) | 0.264 | 47 |  |  |  |
|  | Survival rate (5-p) | 0.310 | 47 |  |  |  |
|  | Larval period (days) | 109482.038 | 424 |  |  |  |
|  | Pupal weight | 28585.921 | 276 |  |  |  |
|  | Female pupa period (days) | 39137.205 | 307 |  |  |  |
|  | Male pupa period (days) | 36656.652 | 301 |  |  |  |
|  | Emergence rate | 0.290 | 47 |  |  |  |
|  | Proportion FA | 0.216 | 47 |  |  |  |
|  | Longevity of female adult | 996.113 | 300 |  |  |  |
|  | Longevity of male adult | 1344.972 | 323 |  |  |  |
|  | Number of eggs produced per female | 2563103.079 | 277 |  |  |  |
|  | Hatching rate | 3.930 | 243 |  |  |  |

The categorical variables were RL (resistant levels of different susceptible or resistant strains) and HaDV2 (infected with HaDV2 or not). Interactions were not significant.

**Supplementary table 5.**
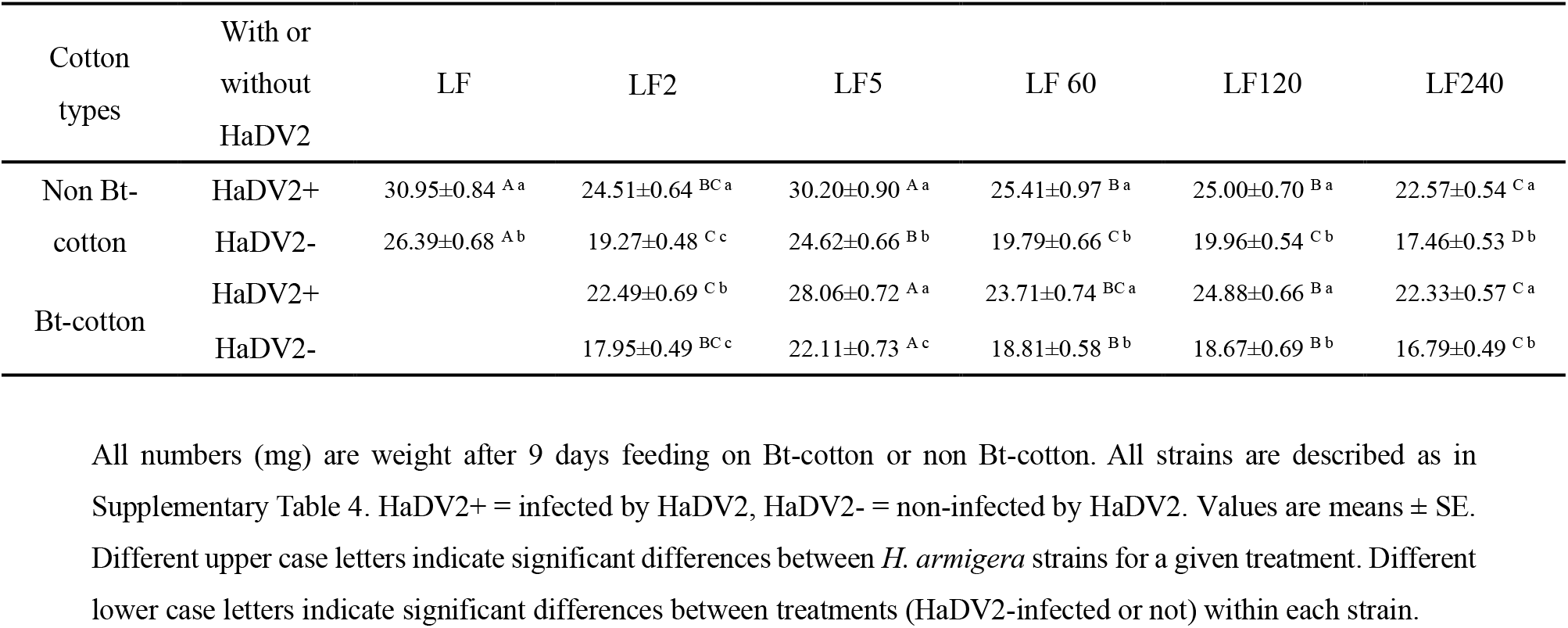
Fitness of the susceptible and resistant strains on Bt- and non-Bt-cotton infected with or without HaDV2.

**Supplementary table 6.** Analysis of variance for weight of cotton bollworm larvae.

| Source | Type III sum of squares | df | Mean square | F | P |
| --- | --- | --- | --- | --- | --- |
| Model | 18650.995 | 77 | 242.221 | 10.762 | 0.000 |
| Intercept | 335764.047 | 1 | 335764.047 | 14918.250 | 0.000 |
| Bt | 415.442 | 1 | 415.442 | 18.458 | 0.000 |
| RL | 7536.653 | 6 | 1256.109 | 55.810 | 0.000 |
| DNV | 7299.219 | 1 | 7299.219 | 324.310 | 0.000 |
| Bt * RL | 120.643 | 4 | 30.161 | 1.340 | 0.253 |
| Bt * HaDV2 | 0.846 | 1 | 0.846 | .038 | 0.846 |
| RL * HaDV2 | 22.977 | 4 | 5.744 | 0.255 | 0.907 |
| Bt * RL * HaDV2 | 28.605 | 4 | 7.151 | .318 | 0.866 |
| Error | 23519.745 | 1045 | 22.507 |  |  |
| Total | 631863.849 | 1123 | 631863.849 |  |  |
| Corrected total | 42170.740 | 1122 | 42170.740 |  |  |

The categorical variables were Bt (feed on Bt-cotton or non-Bt-cotton), RL (resistant levels of different susceptible or resistant *H. armigera* strains) and HaDV2 (infected with DV2 or not). Red numbers mean significant difference. Interactions were not significant.

**Supplementary table 7.**
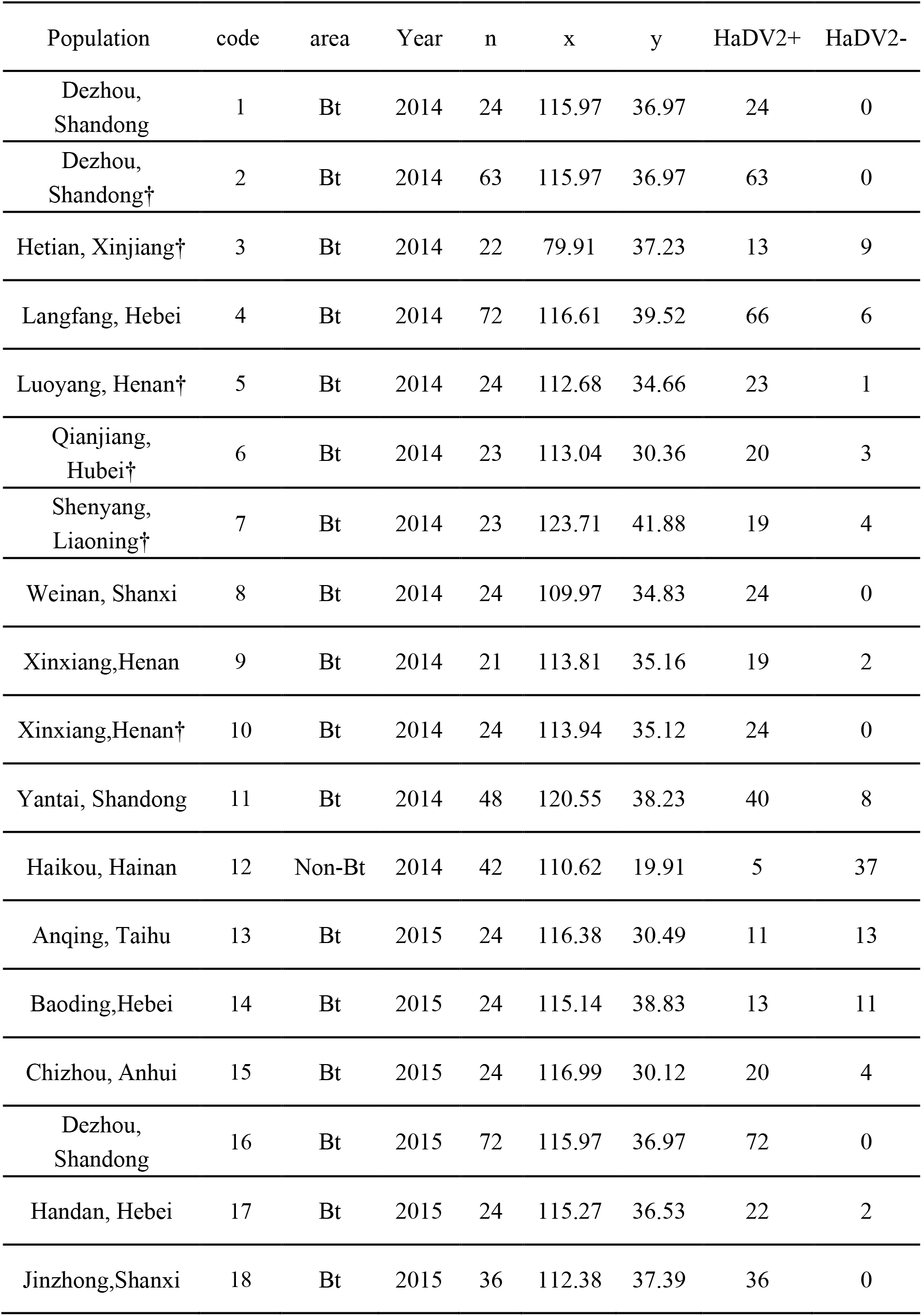
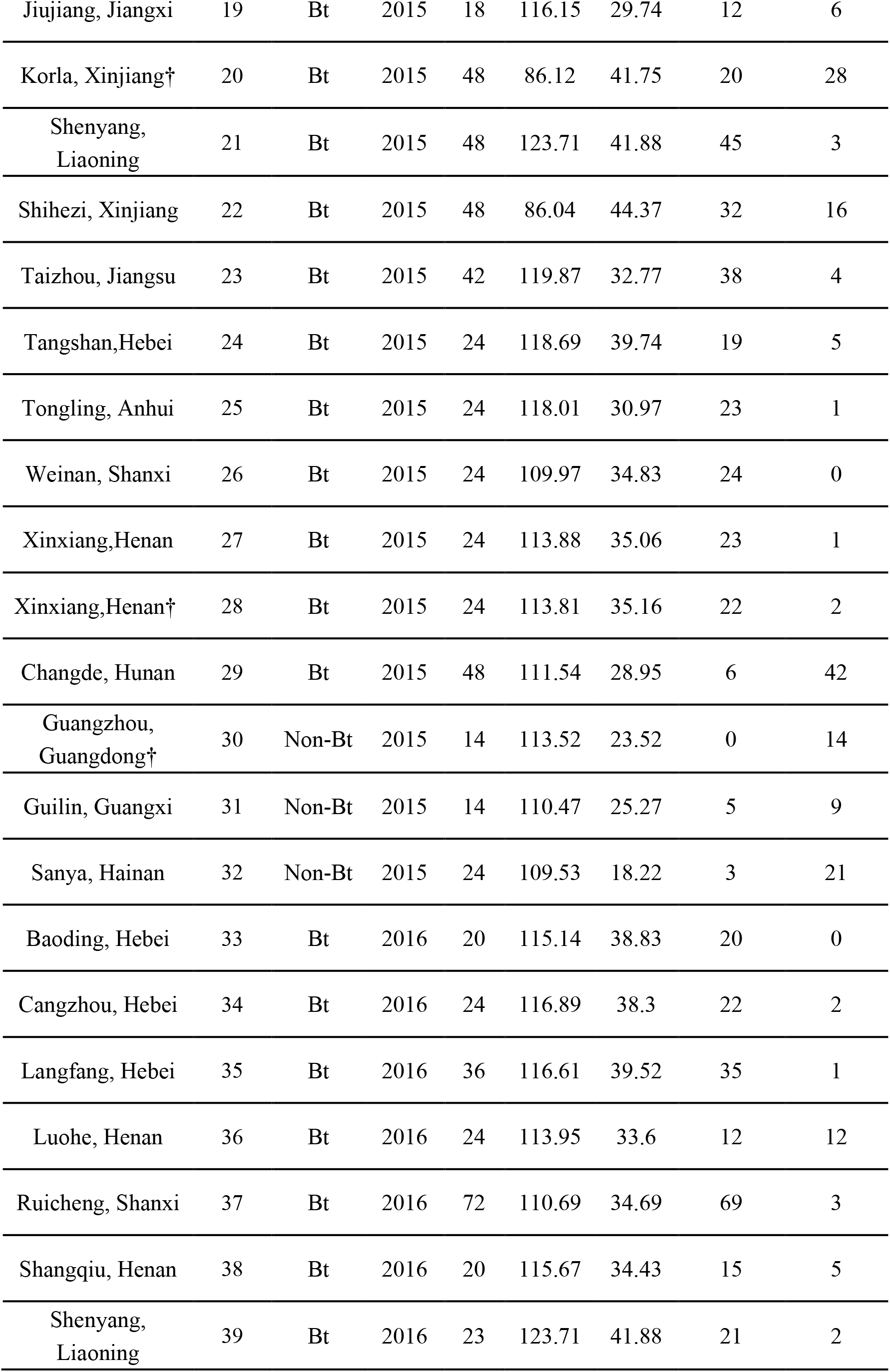
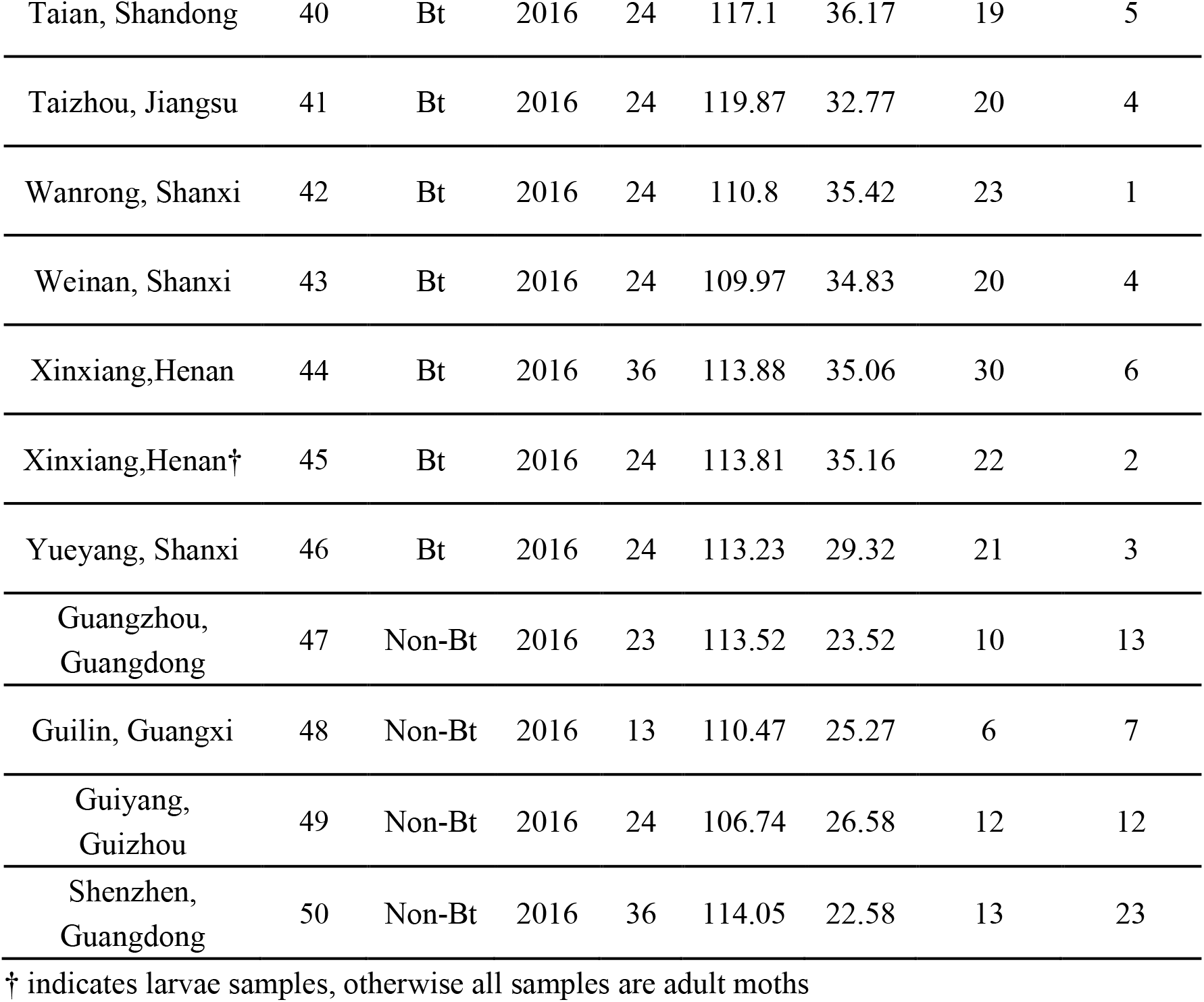
Sample information and infection rates of HaDV2 in the field populations of *Helicoverpa armigera*. See Supplementary Fig. 2 for a map of locations. X: east longitude, Y: northern latitude.

**Supplementary table 8.**
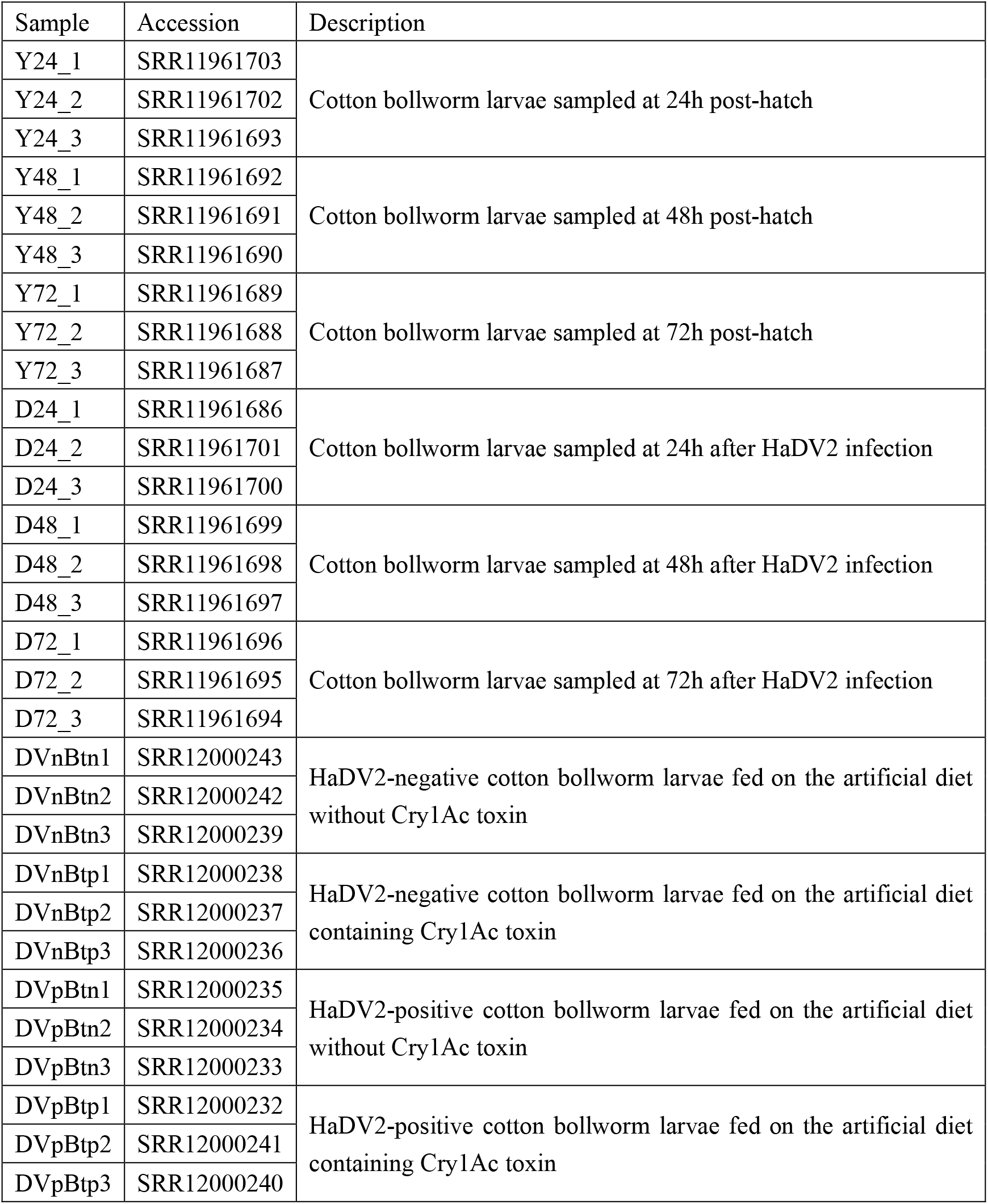
Host information and accessions for samples.

| Sample | Accession | Description |
| --- | --- | --- |
| Y24_1 | SRR11961703 | Cotton bollworm larvae sampled at 24h post-hatch |
| Y24_2 | SRR11961702 |  |
| Y24_3 | SRR11961693 |  |
| Y48_1 | SRR11961692 | Cotton bollworm larvae sampled at 48h post-hatch |
| Y48_2 | SRR11961691 |  |
| Y48_3 | SRR11961690 |  |
| Y72_1 | SRR11961689 | Cotton bollworm larvae sampled at 72h post-hatch |
| Y72_2 | SRR11961688 |  |
| Y72_3 | SRR11961687 |  |
| D24_1 | SRR11961686 | Cotton bollworm larvae sampled at 24h after HaDV2 infection |
| D24_2 | SRR11961701 |  |
| D24_3 | SRR11961700 |  |
| D48_1 | SRR11961699 | Cotton bollworm larvae sampled at 48h after HaDV2 infection |
| D48_2 | SRR11961698 |  |
| D48_3 | SRR11961697 |  |
| D72_1 | SRR11961696 | Cotton bollworm larvae sampled at 72h after HaDV2 infection |
| D72_2 | SRR11961695 |  |
| D72_3 | SRR11961694 |  |
| DVnBtn1 | SRR12000243 | HaDV2-negative cotton bollworm larvae fed on the artificial diet without Cry1Ac toxin |
| DVnBtn2 | SRR12000242 |  |
| DVnBtn3 | SRR12000239 |  |
| DVnBtp1 | SRR12000238 | HaDV2-negative cotton bollworm larvae fed on the artificial diet containing Cry1Ac toxin |
| DVnBtp2 | SRR12000237 |  |
| DVnBtp3 | SRR12000236 |  |
| DVpBtn1 | SRR12000235 | HaDV2-positive cotton bollworm larvae fed on the artificial diet without Cry1Ac toxin |
| DVpBtn2 | SRR12000234 |  |
| DVpBtn3 | SRR12000233 |  |
| DVpBtp1 | SRR12000232 | HaDV2-positive cotton bollworm larvae fed on the artificial diet containing Cry1Ac toxin |
| DVpBtp2 | SRR12000241 |  |
| DVpBtp3 | SRR12000240 |  |

**Supplementary figure 1.**
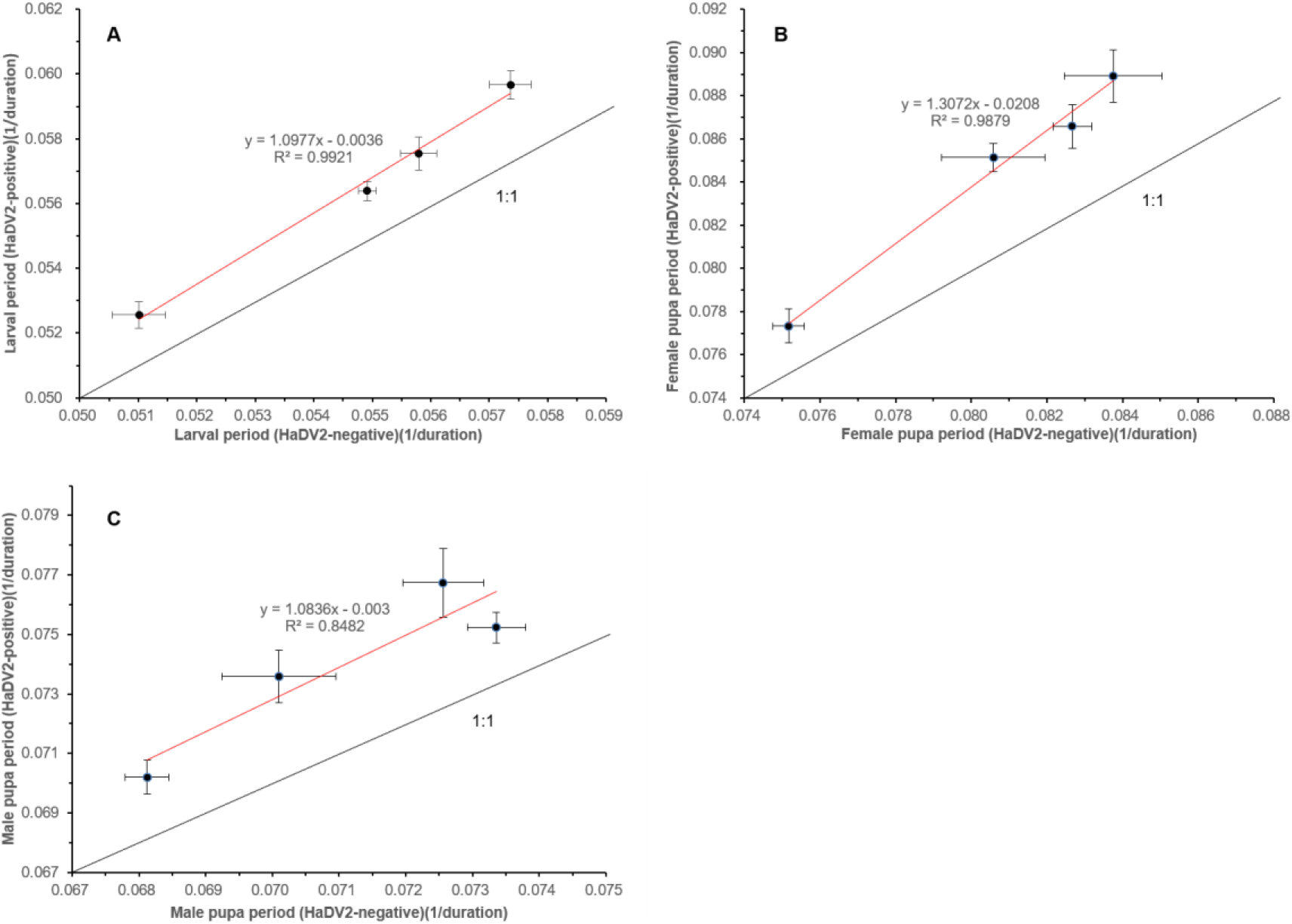
Relationship of different *H. armigera* strains’ larval (A), and pupal (B, C) with or without HaDV2 infection. (A): The x-axis is the larval development rate (1/duration) of different strains (LF, LF5, LF60, LF240) without HaDV2 infection (HaDV2-negative); the y-axis is the larval development rate (1/duration) of different strains (LF, LF5, LF60, LF240) with HaDV2 infection (HaDV2-positive), y = 1.0977x - 0.0036, R² = 0.9921, F = 176.678, df = 1, 2, P = 0.006. (B): The x-axis is the Female pupal development rate (1/duration) of different strains (LF, LF5, LF60, LF240) without HaDV2 infection (HaDV2-negative); the y-axis is the Female pupa period (1/duration) of different strains (LF, LF5, LF60, LF240) with HaDV2 infection (HaDV2-positive), y = 1.3072x – 0.0208, R² = 0.9879, F = 125.211, df = 1, 2, P = 0.008. (C): The x-axis is the Male pupa period (1/duration) of different strains (LF, LF5, LF60, LF240) without HaDV2 infection (HaDV2-negative); the y-axis is the Male pupa period (1/duration) of different strains (LF, LF5, LF60, LF240) with HaDV2 infection (HaDV2-positive), y = 1.0836x - 0.003, R² = 0.8482, F = 7.581, df = 1, 2, P = 0.110.

**Supplementary figure 2.**
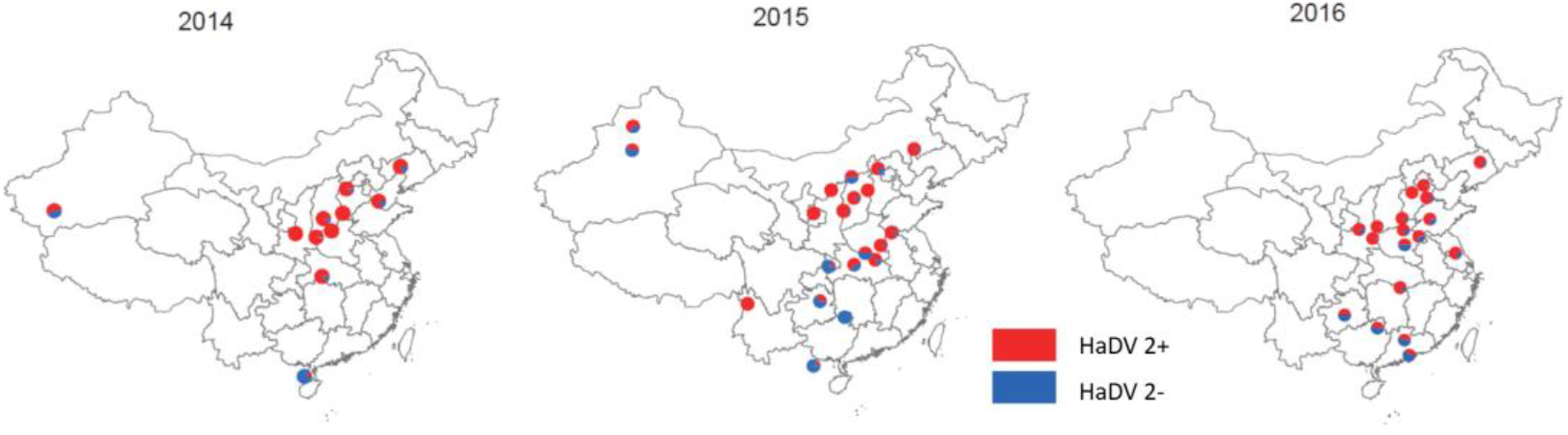
Distribution of HaDV2 in *H. armigera* from different populations. The red proportion of circles refer to infected individuals, and the blue refers to non-infected individuals. There are significant difference of HaDV2 infection rate between the 29 Bt-cotton planting points and 7 non-Bt-cotton planting points (code: 12, 29, 30, 31, 32, 49, 50). The sample information was summarized in Supplementary Table 6.

**Supplementary figure 3.**
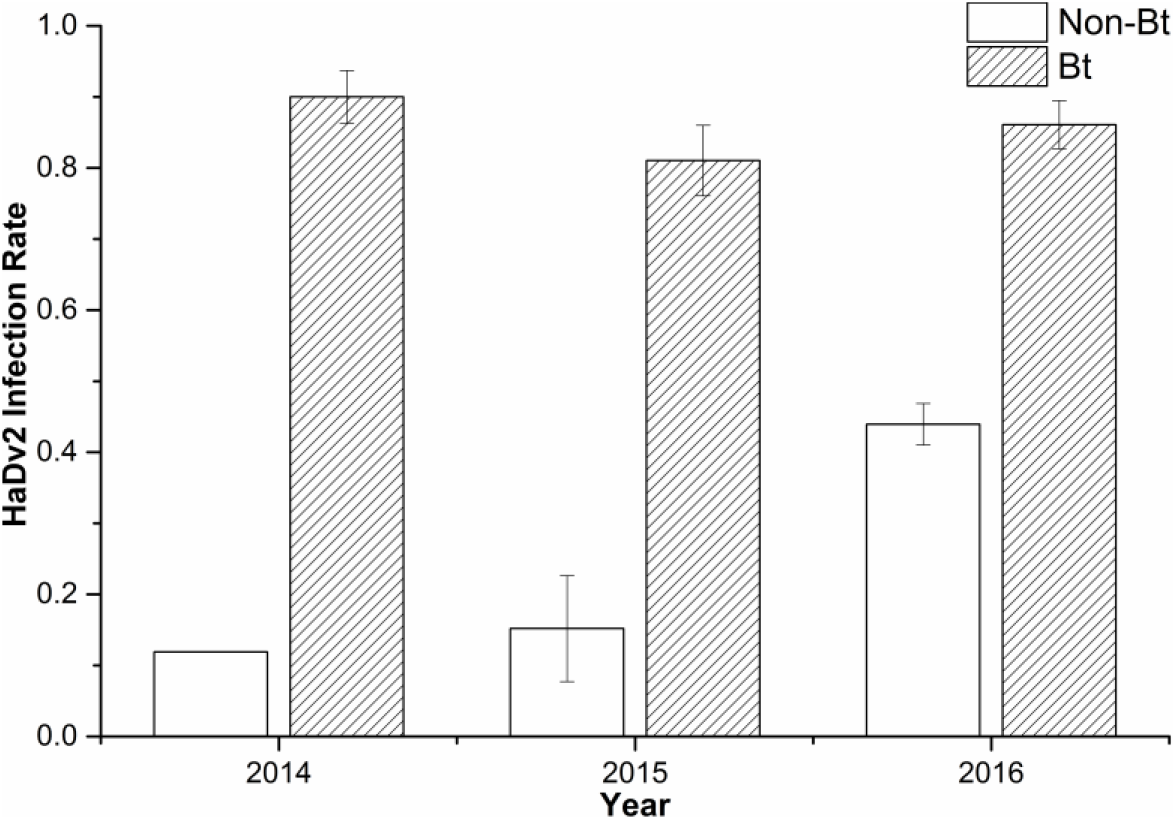
The infection rate of HaDV2 in Bt-cotton and non-Bt-cotton planting areas from 2014 - 2016 (Means ± SE). The detailed information is summarized in Supplementary Table 6 and Supplementary Fig. 2.

**Supplementary figure 4.**
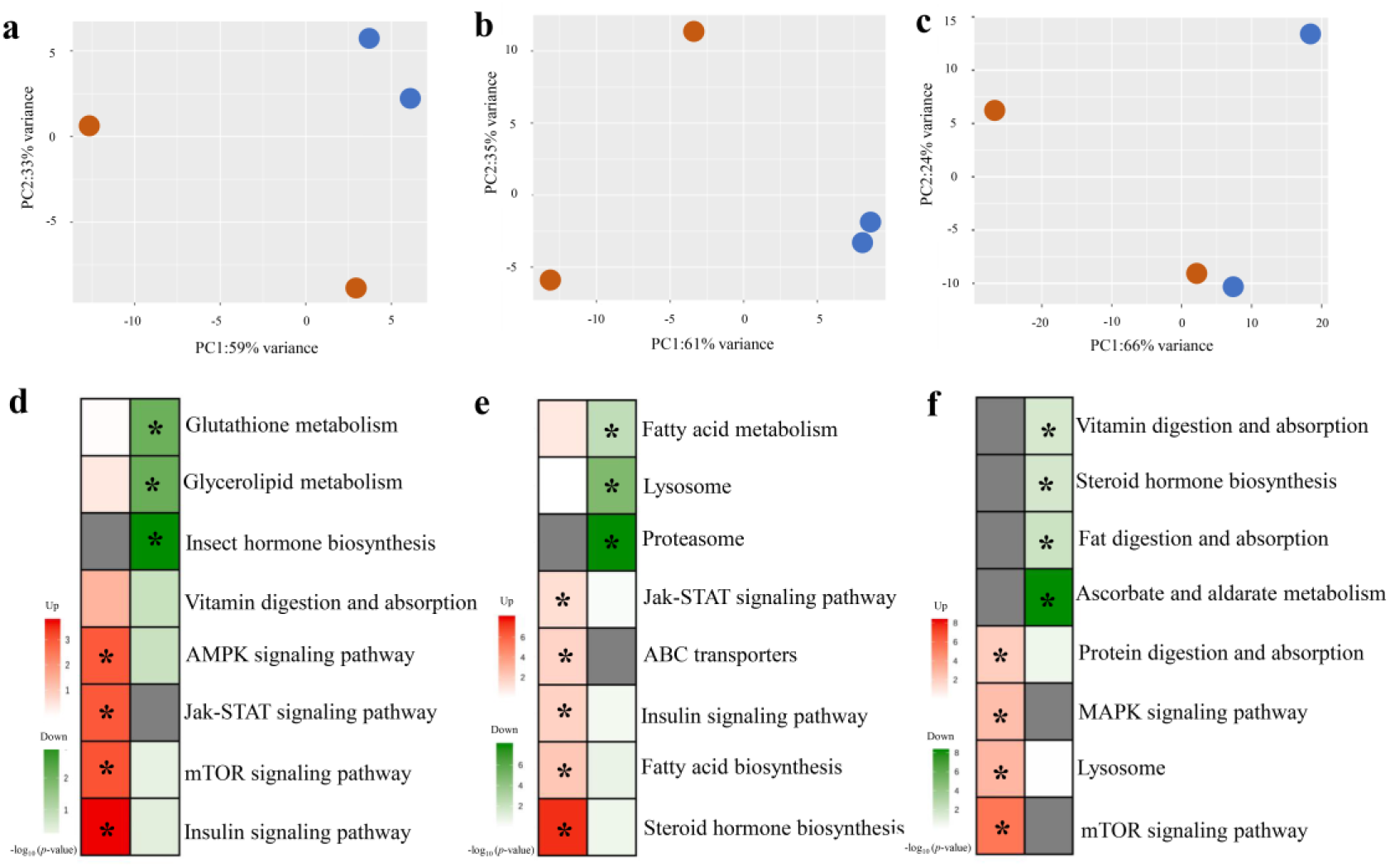
Transcriptome analysis of HaDV2-positive individuals compared to related HaDV2-negative individuals (HaDV2+ vs HaDV2-) of *H. armigera*. (a, b, c) PCA of global gene expression of DEGs at 24 h (a), 48 h (b) and 72 h (c) after HaDV2 inoculation. Blue stands for HaDV2-positive samples and red stands for HaDV2-negative samples. (d, e, f) Heatmaps of –log10 *p*-values of KEGG pathways representing the up- and down-regulated DEGs at 24 h (d), 48 h (e) and 72 h (f). “*” indicate the significantly enriched pathways (p < 0.05). Red color shows up-regulation pathways, green color show down-regulation pathways, gray color shows no value, the redder/greener the color, the lower *P*-values.

**Supplementary figure 5.**
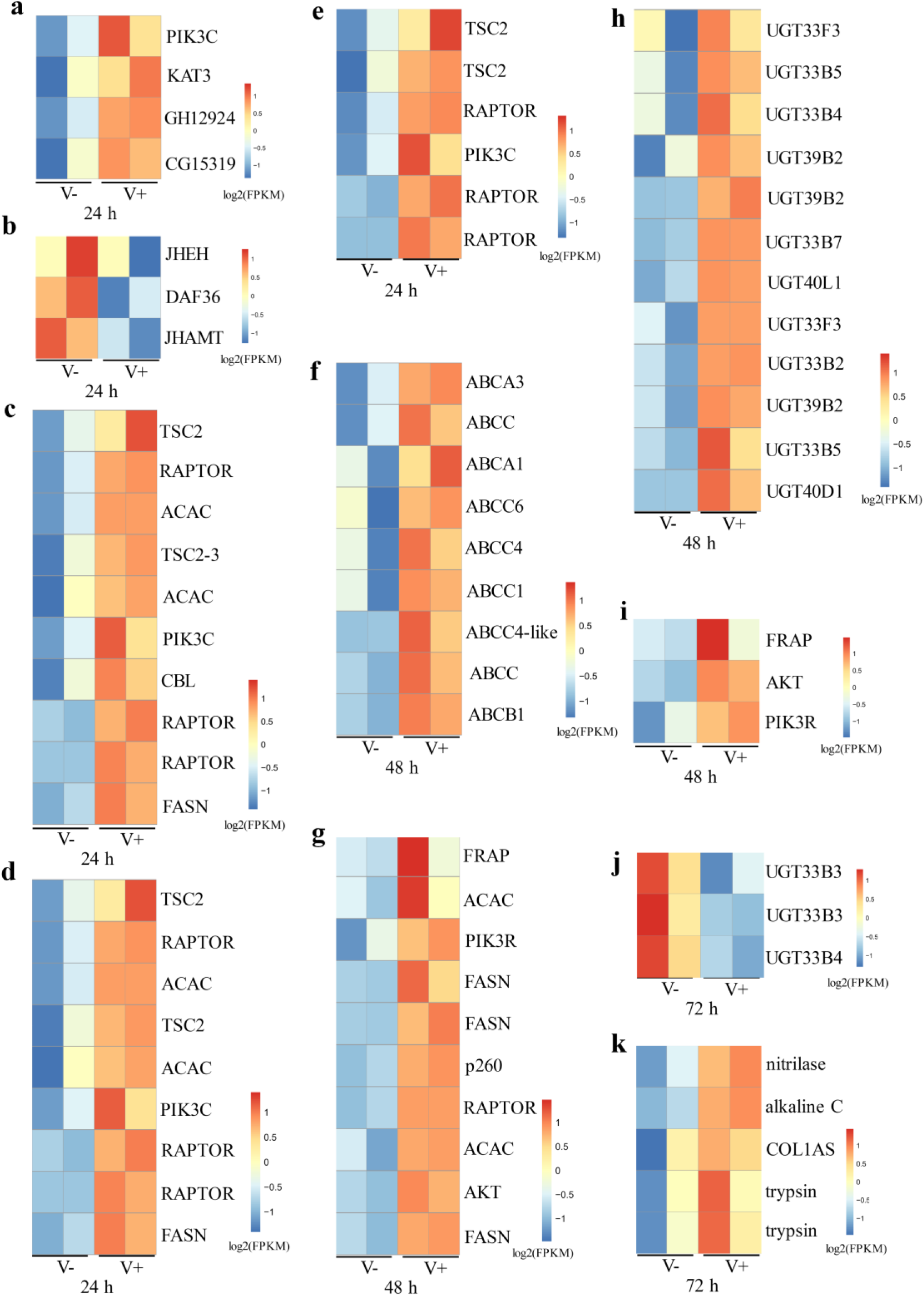
Heatmaps of DEGs related to the expression of significantly enriched pathways of *H. armigera* at 24 h, 48 h and 72 h. The quantity of DEGs with log_2_(FPKM) related to the expression of the Jak-STAT signaling pathway at 24 h (a) and 48 h (i); the insect hormone biosynthesis pathway at 24 h (b); the insulin signaling pathway at 24 h (c) and 48 h (g); the AMPK signaling pathway at 24 h (d); the mTOR signaling pathway at 24 h (e); the ABC transporters at 48 h (f); the steroid hormone biosynthesis pathway at 48 h (h) and 72 h (j); the protein digestion and absorption pathway at 72h (k). V- = HaDV2-negative individuals, V+ = HaDV2-positive individuals. Colors in log2(FPKM) indicate the gene expression levels, the hotter (redder) the color, the higher the gene expression level. The two columns in V- and V+ represents two replications.

**Supplementary figure 6.**
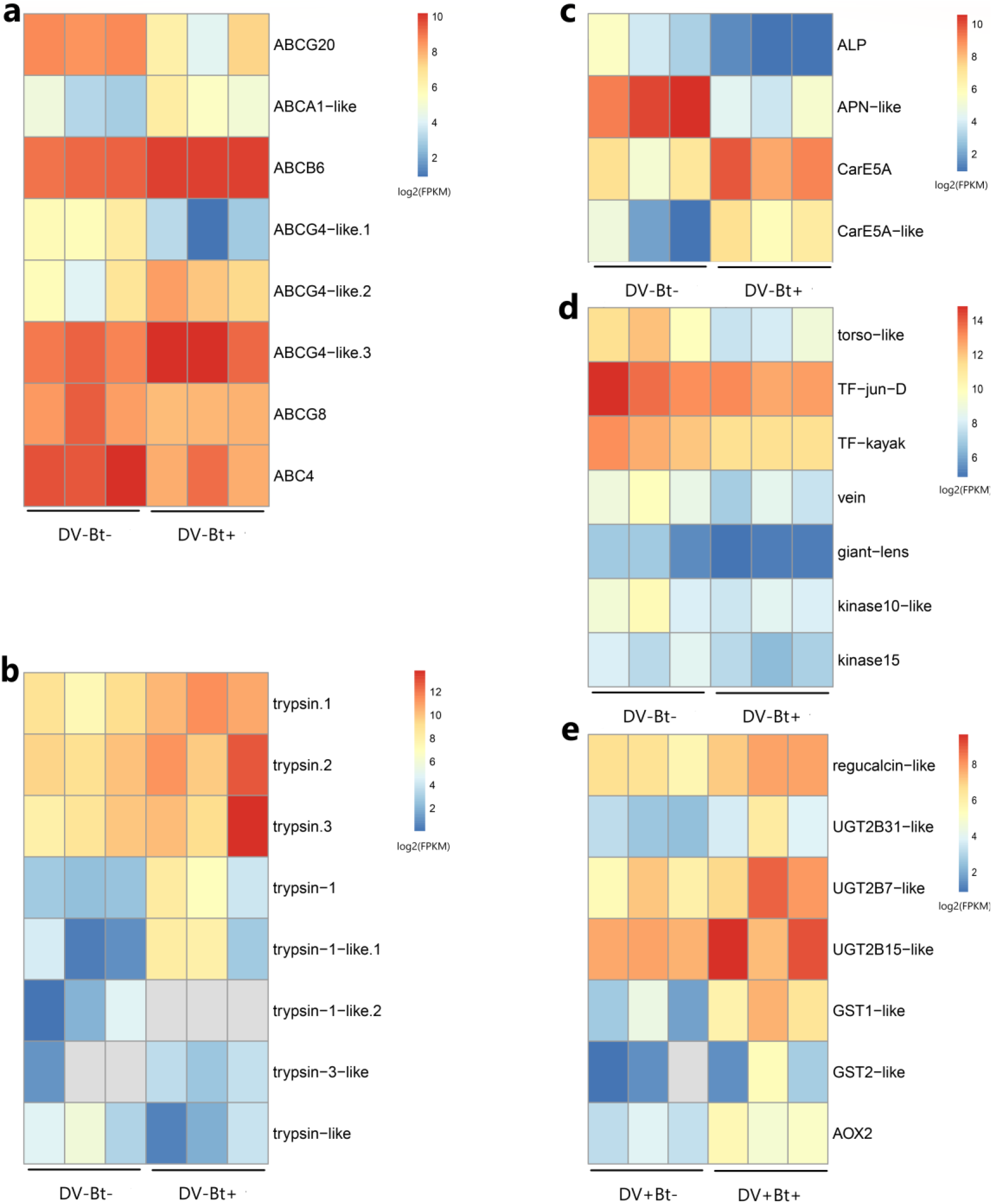
Transcriptome analysis of *H. armigera* after HaDV2 infection and Cry1Ac exposure. The quantity of DEGs with log_2_(FPKM) related to the expression of (a) the ABC transporters; (b) trypsin; (c) Bt receptors and carboxylesterase genes; (d) the MAPK signaling pathway; (e) the drug metabolism pathways. DV- = HaDV2-negative individuals, DV+ = HaDV2-positive individuals. Bt- = larvae were fed on the artificial diet without Cry1Ac, Bt+ = larvae were fed on the artificial diet containing 1 μg/ml Cry1Ac. Colors in log_2_(FPKM) indicate the gene expression levels, the hotter (redder) the color, the higher the gene expression level. The three columns represents three replications.

